# FusedFCR: A Fused Forward Continuation-Ratio model for marker selection along cell-fate trajectories

**DOI:** 10.64898/2026.07.20.739649

**Authors:** Chloe Mattila, Anirban Chakraborty, Peggi Angel, Sha Cao, Kalyani Sonnawane, Elizabeth Hill, Dongjun Chung, Brian Neelon, Souvik Seal

**Affiliations:** Department of Public Health Sciences, Medical University of South Carolina, Charleston, SC, USA; Department of Pharmacology and Immunology, Medical University of South Carolina, Charleston, SC, USA; Department of Biomedical Engineering, Oregon Health & Science University, Portland, OR, USA; Department of Biomedical Informatics, The Ohio State University, Columbus, Ohio, USA

**Keywords:** pseudotime, longitudinal scRNA-seq, cell-fate trajectory, forward continuation ratio, fused lasso, variable selection

## Abstract

Time-course single-cell RNA sequencing (scRNA-seq) data collected across ordered stages provide population-level snapshots of differentiation, disease progression, and aging. Supervised pseudotime methods use observed stage labels to reconstruct continuous progression but generally do not identify marker genes associated with changes from one stage to the next. Unsupervised pseudotime-based marker selection methods infer latent trajectories directly from expression data and identify trajectory-associated genes, but do not explicitly link these associations to the observed stages. We propose FusedFCR, a regularized forward continuation-ratio model that represents cellular progression through a sequence of conditional transitions across ordered stages. FusedFCR combines a lasso penalty for gene selection with a fusion penalty that encourages similar effects across adjacent transitions while allowing transient and direction-changing associations. The resulting transition-specific coefficients support interpretable gene selection and a continuous pseudotime-like projection anchored to the observed developmental stages. In simulations, FusedFCR accurately recovers gene-effect trajectories and improved predictive performance relative to alternative methods. Applied to one mouse and three human datasets (mouse pancreatic beta-cells, human extravillous trophoblast, human induced pluripotent stem cell derived astrocytes, and human endometrial cells during the secretory phase), FusedFCR identifies biologically interpretable genes associated with distinct developmental transitions. Gene set enrichment analysis further reveals stage-specific pathway activity consistent with known developmental biology, while held-out stage-classification accuracy was competitive or superior across both datasets. Together, these results show that FusedFCR complements pseudotemporal ordering by identifying which molecular programs change and when those changes emerge along the developmental trajectory. An accompanying R package is available on GitHub.

## 1 Introduction

Single-cell RNA sequencing (scRNA-seq) has reshaped cell biology by enabling genome-scale molecular profiling of individual cells, revealing cellular heterogeneity, transient cell states, and developmental programs that are obscured in bulk measurements (Linnarsson and Teichmann, 2016). Increasingly, single-cell and related omics technologies are moving beyond static snapshot designs toward longitudinal or time-course profiling, providing temporal information needed to characterize the molecular programs that shape cell-fate trajectories, defined here as ordered transitions across developmental stages (Moris et al., 2016; Rizvi et al., 2017; Ding et al., 2022; Aivazidis et al., 2025; Lee et al., 2025; Furtwängler et al., 2025; Gu et al., 2025; Zhang et al., 2025). For example, Delile et al. (2019) profiled single cells from the cervical and thoracic regions of the developing mouse neural tube between embryonic days 9.5 and 13.5, revealing spatial and temporal gene-expression programs underlying neuronal specification. Similarly, Zhai et al. Zhai et al. (2022) used longitudinal single-cell profiling of paired acute myeloid leukemia (AML) diagnosis–relapse samples to reveal clonal and transcriptional heterogeneity associated with recurrence. These studies highlight the potential of time-resolved scRNA-seq for reconstructing biological progression, while underscoring the need for computational methods that identify interpretable, transition-specific drivers of that progression (Wang et al., 2021).

When observed time-course labels are unavailable, trajectory inference methods use snapshot scRNA-seq data to infer a scale-free pseudotime that orders cells by relative differentiation state and summarizes developmental paths across cell populations (Trapnell et al., 2014; Haghverdi et al., 2016; Cao et al., 2019; Saelens et al., 2019; Wei et al., 2021, 2019; Faure et al., 2023). For time-course scRNA-seq data with observed collection times, however, unsupervised trajectory approaches can be suboptimal because they do not directly incorporate the experimental times at which cells were sampled (Reid and Wernisch, 2016; Weinreb et al., 2018; Tritschler et al., 2019; Ahmed et al., 2019; Lederer and La Manno, 2020). To address this limitation, psupertime (Macnair et al., 2022) introduced a supervised framework that uses collection times as ordinal labels for pseudotime construction, producing latent orderings that better align with known temporal progression than unsupervised methods such as Monocle 2 (Qiu et al., 2017b) and Slingshot (Street et al., 2018). Importantly, psupertime also provides interpretable gene-level effect estimates, enabling the identification of genes broadly associated with developmental progression. More recently, supervised temporal inference methods have been extended using more flexible models, including nonlinear classifiers such as Sceptic (Li et al., 2025) and deep generative models such as StemVAE (Cao et al., 2025). These methods improve the modeling of complex gene-expression patterns for timestamp prediction or temporal-state inference, but place less emphasis on interpretable gene selection. In parallel, identifying marker genes for temporally ordered biological processes remains a central analytical challenge. Existing approaches typically infer pseudotime using trajectory methods and then test each gene independently for dynamic association with the inferred trajectory (Campbell and Yau, 2016; Fischer et al., 2018; Cao et al., 2020; Song and Li, 2021; Fan et al., 2024; Lai et al., 2025). Thus, current methods leave a gap for time-course data: supervised methods use collection times but primarily identify genes associated with overall progression, whereas unsupervised marker-discovery methods detect temporally dynamic genes but usually rely on inferred trajectories rather than directly modeling observed stage-to-stage transitions.

Time-course scRNA-seq data contain ordered sampling times but not classical longitudinal measurements. Because scRNA-seq profiling is typically destructive, each cell is observed only once, so time points represent population-level snapshots rather than repeated measurements of the same cells. This structure naturally motivates ordinal regression frameworks that preserve temporal ordering, particularly continuation-ratio (CR) models, which represent progression as a sequence of conditional transitions (Cox, 1988; Ananth and Kleinbaum, 1997; Archer and Williams, 2012; Suresh et al., 2022; Sun and Cui, 2026). psupertime uses this framework by modeling the ordered time point as a function of gene expression, with each gene assigned a single effect shared across all transitions. It further imposes a lasso (*ℓ*_1_) penalty on these effects to enable gene selection (Tibshirani, 1996). Consequently, a gene is selected based on a single effect estimated across the full sequence of time points, potentially obscuring markers whose association with progression is specific to individual transitions. This is biologically important because genes often show transient or stage-restricted expression along cellular trajectories, making them informative for some transitions but not others (Campbell and Yau, 2016; Fischer et al., 2018; Cao et al., 2020; Song and Li, 2021; Fan et al., 2024; Lai et al., 2025).

Motivated by this gap, we extend the forward continuation-ratio (FCR) model (Seffernick and Archer, 2024) to allow transition-specific gene effects. Let *y* denote a cell’s ordered sampling time and **x** = (*x*_1_*, . . ., x_p_*)*^T^* its *p*-dimensional gene-expression profile. At transition *t*, the model estimates *P* (*y* = *t* | *y* ≥ *t,* **x**), the probability that a cell is observed at time *t* given that it is observed at time *t* or later. This probability is modeled as a function of the linear predictor,logit 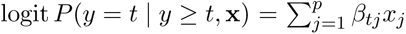. The coefficient *β_tj_* measures the association of gene *j* with this transition: *β_tj_ >* 0 favors observation at time *t*, whereas *β_tj_ <* 0 favors progression to later times. Allowing *β_tj_* to vary across transitions therefore enables the identification of transition-specific marker effects. For parsimonious marker selection, we impose a lasso penalty on the gene coefficients. We additionally impose a one-dimensional fused lasso penalty on successive differences, |*β_tj_* −*β*_(_*_t_*_−1)_*_j_* |, to encourage similar effects across adjacent transitions (Tibshirani et al., 2005; Arnold and Tibshirani, 2016; Adhikari et al., 2019), motivated by evidence that gene expression (e.g., of hallmark genes) evolves gradually across adjacent developmental stages, rather than isolated, independent, stages (Cardoso-Moreira et al., 2019; Ding et al., 2022; Liang et al., 2022). This is not universally true, as for several transient cell population, gene expression shows abrupt shifts Jansky et al. (2021). The fused lasso accommodates both kinds of changes by shrinking adjacent coefficients towards one another only when supported by the data, rather than enforcing uniformity across transitions. Together, these penalties promote gene sparsity and temporal coherence while allowing effects to change when supported by the data. Crucially, such fused lasso-based methods have been utilized in biomarker discoveries for longitudinal sequencing data Nowak et al. (2011); Miah et al. (2025). We call the proposed model FusedFCR and compare it with existing methods in terms of gene-selection performance and prediction accuracy on held-out data across a range of challenging simulation settings. We also develop a new simulation framework tailored to FCR models, addressing the lack of readily available tools for generating data under this model class. We apply FusedFCR to four time-course scRNA-seq datasets: (1) embryonic pancreatic beta-cell development (Qiu et al., 2017a), (2) trophoblast differentiation (Moreno-Irusta et al., 2026), (3) induced pluripotent stem cell derived astrocyte differentiation (Yi et al., 2024), and (4) endometrial transformation during the secretory phase of the menstrual cycle (Wang et al., 2020), identifying pathway-relevant genes with transition-specific effects. A scalable implementation is available as an R package on GitHub.

## 2 Results

### 2.1 Simulation Study

We conducted simulation studies to compare FusedFCR with psupertime in terms of variable-selection performance and to evaluate the predictive accuracy of FusedFCR, psupertime, and Sceptic. Because Sceptic does not provide gene-level coefficient estimates, it was included only in the predictive accuracy comparison. Each approach was assessed across settings with increasing dimensionality (*p*) relative to the number of cells (*N*). We considered two simulation scenarios that differ in how the gene expression data and true coefficients are structured. In both scenarios, the outcome *y_i_* is generated from the FCR model (Equations 4.3 and 4.4). In the *non-temporal* scenario, gene expression values are generated independently from Gaussian distributions, without reference to an underlying developmental process. In the *temporal* scenario, the expression levels of signal genes are generated as functions of latent developmental time, which also determines the transition probabilities. Thus, gene expression and stage progression share a common latent structure.

For both scenarios, we considered *T* = 6 ordered stages with five transitions (i.e., 1 → 2 → 3 → 4 → 5 → 6). Outcomes were generated sequentially (Eq. 4.1) and all cells entered into the risk set for the first transition, and at each transition, a Bernoulli random variable was drawn using the corresponding transition probability. If observation *i* stayed at transition *t*, then *y_i_* = *t* and it was removed from subsequent risk sets; observations that continued through all five transitions were assigned *y_i_* = 6. However, readers should note that simulating realistic data from the FCR model is nontrivial, as the intercept vector ***α***_0_ jointly governs label balance and prediction accuracy under the true model, and improving one property can come at the cost of the other. We address this by choosing ***α***_0_ differently across our two scenarios to target distinct, complementary properties, favoring balanced labels in one case and higher true-model accuracy in the other, allowing us to evaluate coefficient recovery and predictive performance under the settings best suited to each. Additional details are provided in the Supplementary Material Section S1.

#### 2.1.1 Non-temporal simulation scenario

In the non-temporal scenario, gene expression is drawn independently as *x_ij_* ∼ N (0, 1) and does not vary temporally. Six genes have non-zero coefficients, with two patterns (Table 1):

1. *fixed* signal genes maintain a positive or negative effect across all five transitions, and
2. *varying* signal genes have zero effect at early transitions and a positive effect at later transitions, reflecting a delayed activation, later stage marker gene.

**Table 1:** True signal structure for non-temporal simulations.

| Gene | Expression pattern | True coefficient trajectory |
| --- | --- | --- |
| Gene 1 | Fixed | (1.0, 1.0, 1.0, 1.0, 1.0) |
| Gene 2 | Varying | (0.0, 0.0, 1.0, 1.0, 1.0) |
| Gene 3 | Fixed | (-1.0, -1.0, -1.0, -1.0, -1.0) |
| Gene 4-6 | Varying | (0.0, 0.0, 0.0, 1.0, 1.0) |

The remaining (*p* − 6) genes have a zero coefficient at each transition. The intercept vector was set to ***α***_0_ = (−2.0, −1.5, −1.2, 0.0, 2.0), chosen to produce approximately balanced observation counts across the six outcome categories.

Figure 1A compared the estimated coefficient distributions across 50 replicates under *N* = 1000*, p* = 20 with three nonzero genes (two fixed, one varying; Table 1 gene 1-3) with ***α***_0_ = (−2.0, −1.5, −1.0, −0.5, 1.2), chosen for balanced label distribution. With few non-signal genes competing for selection, FusedFCR’s estimates were close to the truth across all transitions. psupertime performed poorly despite the favorable signal environment, where for the positive *fixed* effect gene, the median estimate was close to the truth but was highly variable; for the negative effect gene the interquartile range was tighter but fell entirely in the wrong direction. For the *varying* effect gene, psupertime’s median estimate was in the wrong direction. We focus on coefficient recovery rather than predictive accuracy as the primary evaluation in this scenario, because in this setup all the methods achieved similar prediction accuracy.

**Figure 1:**
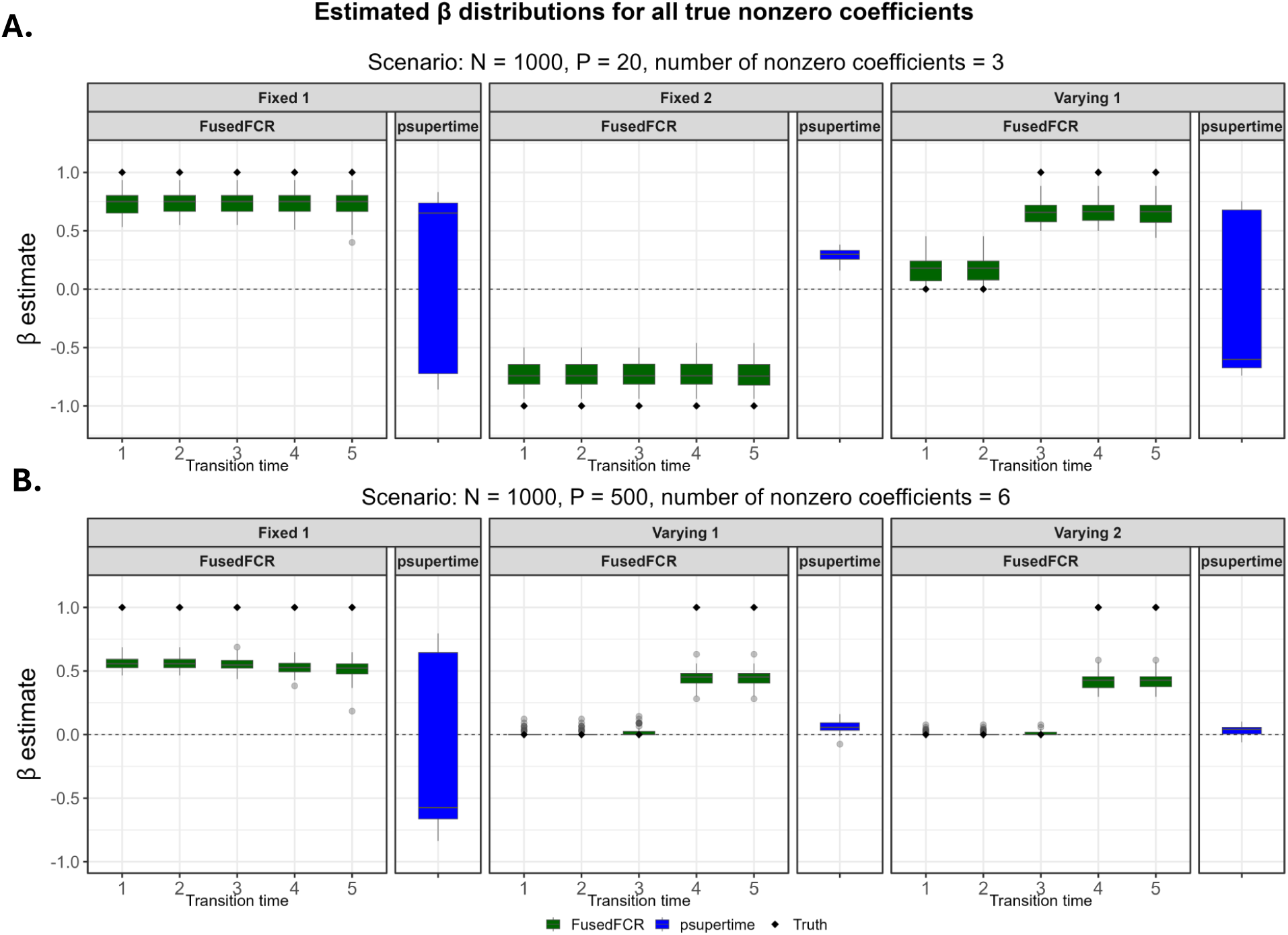
Estimated coefficient distributions across simulation replicates under two non-temporal simulation scenarios. (A) *N* = 1000, *p* = 20, 3 non-zero genes. Signal structure follows the first three genes of Table 1. Each panel shows results for one signal type (Fixed 1, Fixed 2, Varying). (B) *N* = 1000, *p* = 500, 6 non-zero genes; three of the six genes with true non-zero coefficients are shown, one per signal type (Fixed, Varying 1, Varying 2). In both panels, for FusedFCR (green), transition-specific estimates are shown at each of the five transitions; for psupertime (blue), a single pooled estimate is shown. Black diamonds indicate the true coefficient value at each transition.

Figure 1B repeated the comparisons in a higher dimension, *N* = 1000 and *p* = 500, for three of the six nonzero genes (one fixed, two varying signals; Table 1). FusedFCR recovered the transition-specific estimates that are concentrated around the truth, with modest underestimation toward zero, as expected under *ℓ*_1_ penalization (Szendrei et al., 2024) and correctly captured both the early transition null effects and the positive later effects. psupertime, constrained to a single estimate per gene, performed poorly on both signal types. For the *fixed* signal genes, despite this being the setting best aligned with psupertime’s single-coefficient assumption, estimates were highly variable and centered near zero. For the *varying* signal genes, psupertime is not only structurally unable to properly detect the delayed activation signals, but also barely picked up any signal at all. Together, with the lower dimension simulation above, these results indicate that psupertime’s instability and structural blindness to varying effect genes are not artifacts of high dimensionality, but direct consequences of its single coefficient parameterization.

#### 2.1.2 Temporal simulation scenario

For each temporal scenario, we generated a predictor matrix **x** ∈ R*^N^*^×^*^p^*, representing gene expression measurements for *p* genes across *N* observations. As in the nontemporal scenario, each gene’s expression was first drawn independently across genes from N (0, 0.2^2^). Unlike in the preceding scenario, the expression levels of the first three genes were generated as smooth functions of the underlying developmental time, with Gaussian background variation added (see the Supplementary Material Section S1). Their expression therefore evolves continuously along the developmental trajectory rather than changing abruptly between discrete states.

The true coefficient matrix ***β*** ∈ R*^p^*^×5^ was sparse, with only the first three genes carrying non-zero coefficients, so that here both the gene expression distributions and the true regression coefficients vary over developmental time, rather than only the true regression coefficients, as in the nontemporal case. Their transition-specific coefficient signal trajectories are shown in Table 2, and reflect early, middle, and late-stage expression patterns, respectively. For all *t >* 3, ***β****_tj_* = **0**. The intercept vector was set to ***α***_0_ = (−4.5, −4.2, −4.0, −3.2, −2.0). These values were chosen to produce non-extreme transition probabilities and to avoid concentrating observations to the earliest or final outcome category.

**Table 2:** True signal structure for temporal simulations.

| Gene | Expression pattern | True coefficient trajectory |
| --- | --- | --- |
| Gene 1 | Early marker | (5.5, 3.0, 0.0, -3.0, -5.5) |
| Gene 2 | Middle/transient marker | (-2.5, 2.0, 5.5, 2.0, -2.5) |
| Gene 3 | Late marker | (-5.5, -3.0, 0.0, 3.0, 5.5) |

In these temporal simulations, both methods tend to produce many near-zero estimates, we applied simple effect-size thresholding to retain the most strongly selected genes and avoid counting negligible effects as selected. Specifically, we considered three thresholds for the absolute estimated effect size,|β^| > *δ*, with *δ* ∈ {0, 0.05, 0.10}, to evaluate prediction accuracy with respect to the high confidence selected genes. Any coefficient estimate below the applied threshold in absolute value was treated as null and excluded from the selected genes for both psupertime and our FusedFCR at either the global or transition-specific level, respectively. Sceptic does not provide coefficient estimates so no thresholding was done.

Figure 2 summarizes the prediction accuracy on the 20% held-out test set across the two temporal simulation settings at the three *δ* thresholds. In the *N* = 1000 and *p* = 20 setting, all three methods perform similarly, with FusedFCR and Sceptic achieving similarly high prediction accuracy over the 50 replicates while psupertime trailed by 11 percentage points. With increased dimensionality (*p* = 500), a similar pattern held, where FusedFCR and Sceptic again performed similarly, with Sceptic taking a small lead, while psupertime remained lowest, about 12 percentage points below FusedFCR. In this setting, coefficient thresholding did not affect the prediction accuracy of either psupertime or FusedFCR, although its impact became highly apparent in the real-data analyses (see Section 2.2).

**Figure 2:**
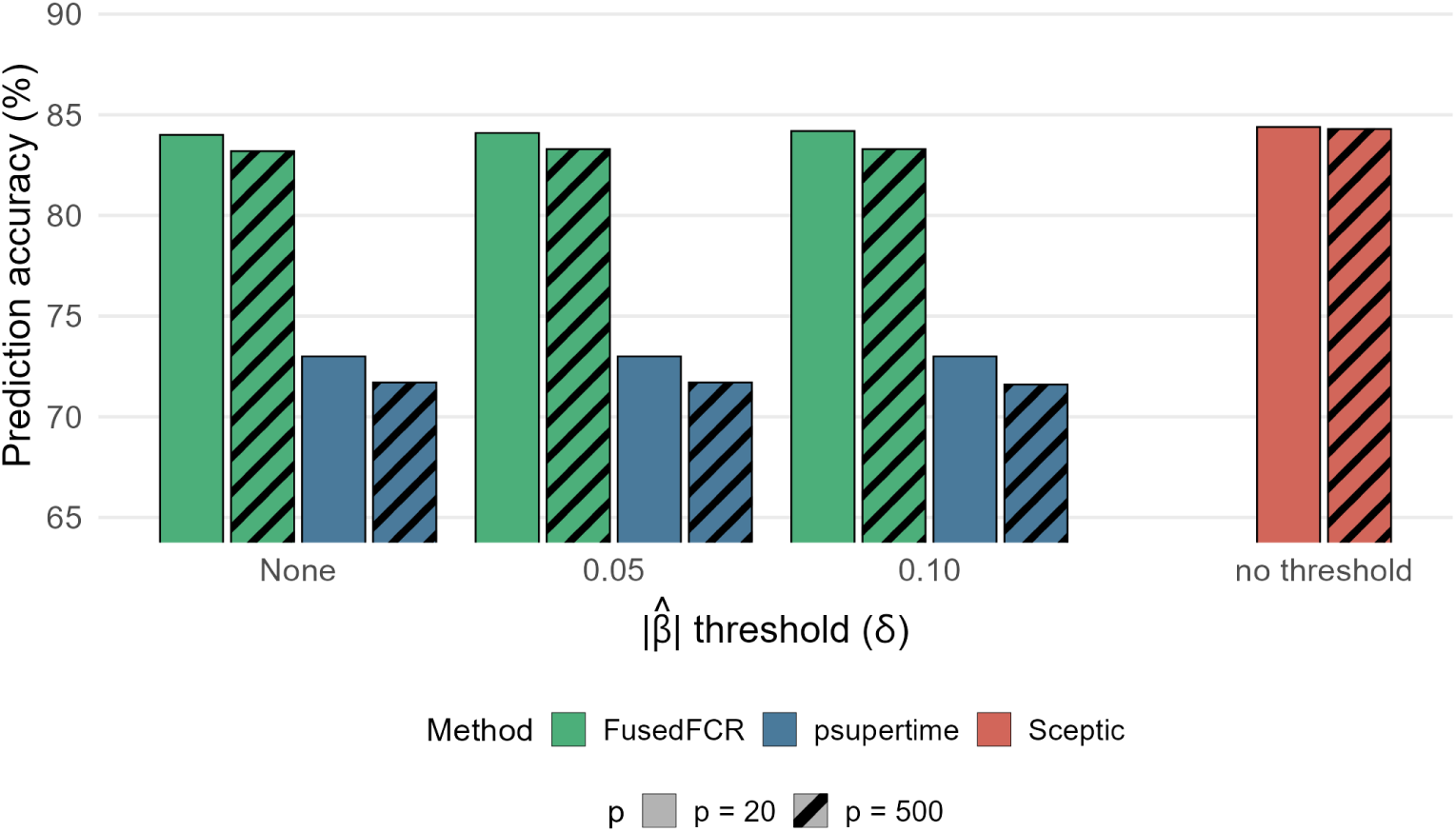
Prediction accuracy (%) on held-out test data from temporal simulation scenarios (rounded to two decimal points), reported as mean *±* standard deviation across 50 replicates. Two simulation settings are shown here (*N* = 1000, *P* = 20 and *N* = 1000, *P* = 500), each with 3 true nonzero predictors per transition.

### 2.2 Real Data Analysis

We analyzed three scRNA-seq datasets that differ in the number of cells and outcome labels or stages (Table 3). For both datasets, we applied the same preprocessing, quality control, and filtering steps, as outlined below.

**Table 3:**
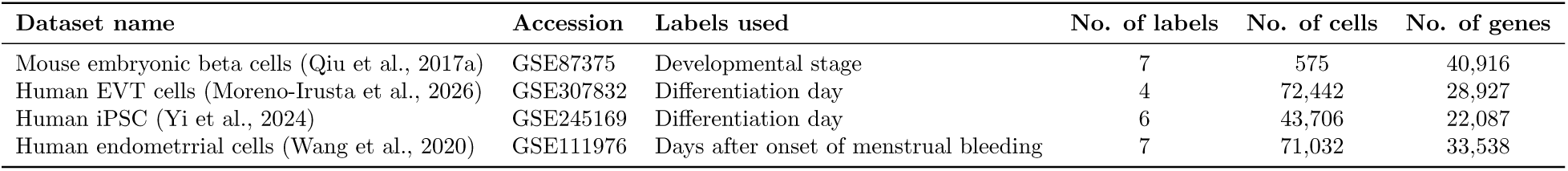
Details of datasets used in benchmarking of cell ordering accuracy.

| Dataset name | Accession | Labels used | No. of labels | No. of cells | No. of genes |
| --- | --- | --- | --- | --- | --- |
| Mouse embryonic beta cells (Qiu et al., 2017a) | GSE87375 | Developmental stage | 7 | 575 | 40,916 |
| Human EVT cells (Moreno-Irusta et al., 2026) | GSE307832 | Differentiation day | 4 | 72,442 | 28,927 |
| Human iPSC (Yi et al., 2024) | GSE245169 | Differentiation day | 6 | 43,706 | 22,087 |
| Human endometrial cells (Wang et al., 2020) | GSE111976 | Days after onset of menstrual bleeding | 7 | 71,032 | 33,538 |

#### 2.2.1 Data Preprocessing and Quality Control

We applied standard scRNA-seq preprocessing steps, including removal of mitochondrial genes, low-quality cells, and low-count genes. We then normalized the data using scuttle (McCarthy et al., 2017) and selected the top 2,000 highly variable genes (HVGs) using scran (Lun et al., 2016).

###### Quality control and preprocessing pipeline *(applied per dataset from Table 3)*

1: **Quality metrics**: Compute per-cell library size, number of detected genes, and mitochondrial percentage (McCarthy et al., 2017). Identify mitochondrial genes by an MT-/mt- prefix and exclude them from analysis.

2: **Low-quality cell removal:** Using developmental-stage label as the batching variable, flag and remove cells with abnormally low library size, low detected gene count, or high mitochondrial percentage (MAD > 3) relative to other cells of the same label; outlier thresholds are computed separately within each label.

3: **Gene filtering:** Retain genes detected (count > 0) in at least 10 cells.

4: **Normalization and HVG selection:** Log-normalize the expression data and model gene-level mean–variance trends. Select the top 2,000 genes by biological variance as HVGs.

#### 2.2.2 Data Curation and Selection Criterion

Gene co-expression in scRNA-seq data induces multicollinearity where correlated genes carry overlapping information, making their proper selection challenging and can undermine gene-level inference (Crow and Gillis, 2018). To address this, we removed genes contributing to this excess multicollinearity prior to model fitting. Pairwise Pearson correlations among the HVGs were computed using the caret package (Kuhn and Max, 2008) based on the log-normalized (cells × genes) matrix. Flagged genes were removed so that pairwise correlations among all remaining genes stay below a chosen cutoff. Two such cutoffs were considered: 0.75 (Case A) and 0.50 (Case B); Case A retains more genes and is presented as the primary analysis. To evaluate predictive performance, 80% of the data was randomly allocated for training with the remaining 20% reserved for testing. As in the temporal simulation study, we applied post-hoc inclusion thresholds to the absolute value of each coefficient estimate |β^| > *δ* and assessed prediction accuracy using the genes retained at each threshold. Sceptic does not provide coefficient estimates so no thresholding was done.

#### 2.2.3 Mouse Embryonic Beta Cells

We analyzed a pancreatic beta cell dataset (Qiu et al., 2017a) from mice sequenced at various developmental stages, embryonic (E17.5), neonatal (P0, P3, and P9), juvenile (P15 and P18), and adult (P60). There are 40,916 genes and 575 beta cells. After quality control filtering (Section 2.2.1) within each developmental stage, 571 cells remained. Case A retained *p* = 1,939 genes, Case B retained *p* = 1,857 genes. The data were split into a training set (80%, *n* = 459) and a held-out test set (20%, *n* = 112). FusedFCR, psupertime, and Sceptic were fit on the training data and evaluated on held-out cells, with performance assessed using prediction accuracy, defined as the proportion of test-set cells whose predicted stage matched their observed stage. For FusedFCR and psupertime, gene-selection performance was further evaluated using individual effect estimates and simple thresholding as described in the temporal simulations (Section 2.1.2). A central feature of FusedFCR is the fused penalty on adjacent transition-specific coefficients, *β_tj_*, which shrinks a gene’s estimate at one transition toward its estimate at a neighboring transition unless the data support a genuine change. This allows sustained effects to be recovered as smooth trajectories, while still allowing for sharper transition specific effects or sign reversals when warranted, a flexibility not possible in a single-coefficient model like psupertime.

In the Case A test data, FusedFCR significantly outperformed both psupertime and Sceptic across all coefficient thresholds, |β^| > *δ* (Fig. 3A). As the threshold increased from *δ* = 0 to *δ* = 0.1, FusedFCR retained most of its predictive accuracy, with accuracy decreasing by only 6.2% despite a 62.9% reduction in the number of selected genes. This suggests that its larger-magnitude coefficients captured most of the predictive signal. In contrast, psupertime accuracy decreased by 21.5% following a comparable 53.5% reduction in selected genes, indicating that many of its smaller-magnitude coefficients contributed meaningfully to prediction. These findings suggest that coefficient magnitude may provide a less reliable measure of gene importance for psupertime, with potential implications for downstream interpretation, including gene set enrichment analysis. A similar pattern was observed in Case B, where accuracy declined by only 9.0% for FusedFCR, compared with 36.6% for psupertime across the considered thresholds.

**Figure 3:**
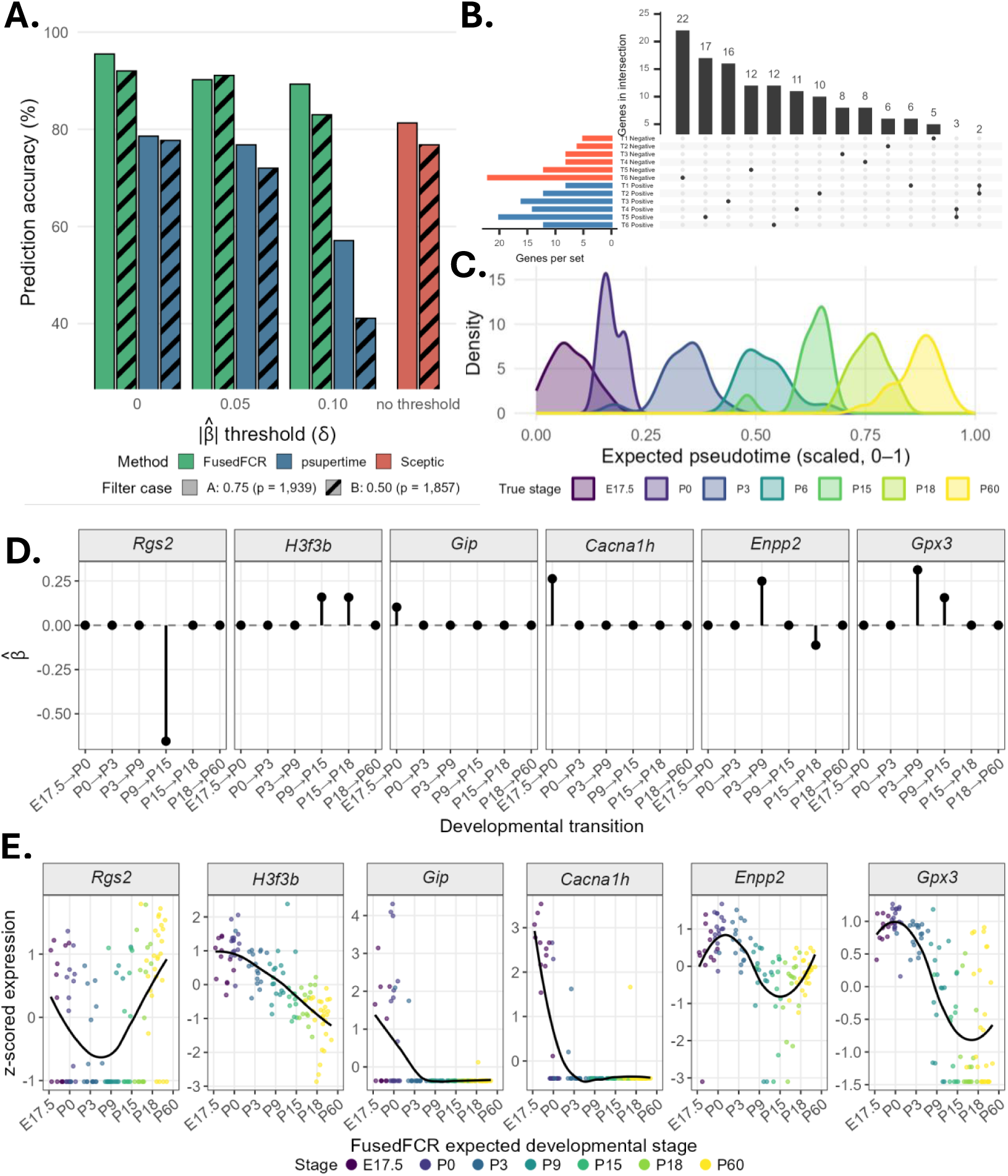
**A.** Gene selection prediction accuracy (%) by method and correlation-filtering threshold in the mouse beta cell dataset (GSE87375) on the held-out test set. **B.** UpSet plot of genes selected by FusedFCR across the six developmental transitions (*|β̂| >* 0.1), colored by coefficient sign (positive, blue; negative, red). **C.** Density plot of FusedFCR pseudotime in the held-out test set (Case A, *δ* = 0.10). **D.** Estimated transition-specific FusedFCR coefficients for six representative genes. **E.** Z-scored log expression for the same genes against FusedFCR expected developmental stage, with LOESS-smoothed trends.

Across the six transitions, for Case A with *δ* = 0.10, few, 12, genes were selected at two or more transitions while 133 genes were selected at a single transition (Fig. 3B). Fig. 3D-E shows, for six highlighted genes, the estimated FusedFCR coefficients at each transition (Fig. 3D) alongside z-scored expression across the FusedFCR estimated pseudotime (Fig. 3E). Beta cell proliferation is highest around the neonatal stages and declines through adulthood (Lee et al., 2021b) and *H3f3b*, which aids in the proliferation of beta cells by encoding the histone H3.3 (Stekelenburg et al., 2022) shows a corresponding decline through the developmental stages (Fig. 3E). FusedFCR positively selected *H3f3b* at two consecutive later transitions (P9 → P15 and P15 → P18), reflects the fused penalty borrowing strength across adjacent estimates to recover the persistent decline in expression. *Gip* and *Cacna1h* were each selected with a single positive effect confined to the first transition (*β̂*_1_*_,Gip_*= 0.10 and *β̂*_1_*_,Cacna1h_*= 0.26), consistent with their high E17.5 expression falling to a low flat postnatal plateau and *Gip*’s role in pancreatic differentiation in embryo (Prasadan et al., 2011). *Enpp2*, which promotes beta cell proliferation and inhibits apoptosis (Zhou et al., 2026), showed a positive effect at the third transition (P3 → P9; *β̂*_3_*_,Enpp2_* = 0.25) followed by a negative effect at the fifth (P15 → P18; *β̂*_5_*_,Enpp2_* = −0.11), mirroring an expression profile that rises to a P3-P9 peak before declining and the sign reversal tracks the rise and then fall shape directly. *Gpx3* showed positive effects at the third and fourth transitions (*β̂*_3_*_,Gpx3_*= 0.31 and *β̂*_4_*_,Gpx3_*= 0.16), consistent with the expression peaking across P3 to P9 before gradually declining. Finally, *Rgs2* carried the single largest-magnitude effect in the panel, a negative coefficient at the fourth transition (P9 → P15; *β̂*_4_*_,Rgs2_* = −0.65), matching the U-shaped expression profile with a trough centered on that transition. This illustrates FusedFCR’s ability to localize a single dominant transition within an otherwise non-monotonic trajectory, and more broadly demonstrate that the fused penalty is data-adaptive. Transition and direction-specific GSEA identified cAMP-mediated signaling as the predominant enriched program at the first transition (KEGG mmu04024; FDR *p* = 7.99 × 10^−5^), driven by *Npy*, *Crhr2*, *Gip*, and *Sox9*, consistent with cAMP’s role as a central second messenger in beta cell fate and proliferation (Tengholm and Gylfe, 2017) (full results in Supplemental Material Section S2).

#### 2.2.4 Human Extravillous Trophoblast Differentiation

We next analyzed a scRNA-seq dataset on extravillous trophoblast (EVT) differentiation (Moreno-Irusta et al., 2026), which profiled human trophoblast stem (TS) cells at differentiation day 0 (D0) and EVT cells at three subsequent time points (D3, D6, and D8). The dataset comprised *n* = 72,442 cells across the four time points. As in the preceding analysis, we considered two feature sets: *p* = 1,982 HVGs in Case A and *p* = 1,781 in Case B. We then applied FusedFCR, psupertime, and Sceptic to both cases.

The data were divided into a training set (80%, *n* = 57,995) and a held-out test set (20%, *n* = 14,487), with all methods trained on the former and evaluated on the latter. In Case A, FusedFCR achieved a test-set prediction accuracy of 97.4%, compared with 90.4% for psupertime and 99.0% for Sceptic (Fig. 4A). As in the beta-cell analysis, increasing the coefficient threshold reduced both the number of selected genes and prediction accuracy for FusedFCR and psupertime. However, the loss in accuracy was modest relative to the reduction in gene count. For FusedFCR, the number of uniquely selected genes decreased by 85.6%, while accuracy declined by only 1.8%. For psupertime, a comparable 88.2% reduction in gene count was accompanied by a larger 5.4% decline in accuracy. The contrast was more pronounced in Case B: FusedFCR reduced the number of selected genes by 84.6%, from 1,766 to 272, with only a 2.2% decline in accuracy, whereas psupertime reduced the gene count by 88.7% but exhibited a substantially larger 20.1% decline in accuracy. This pattern was consistent with the findings from the beta-cell analysis.

**Figure 4:**
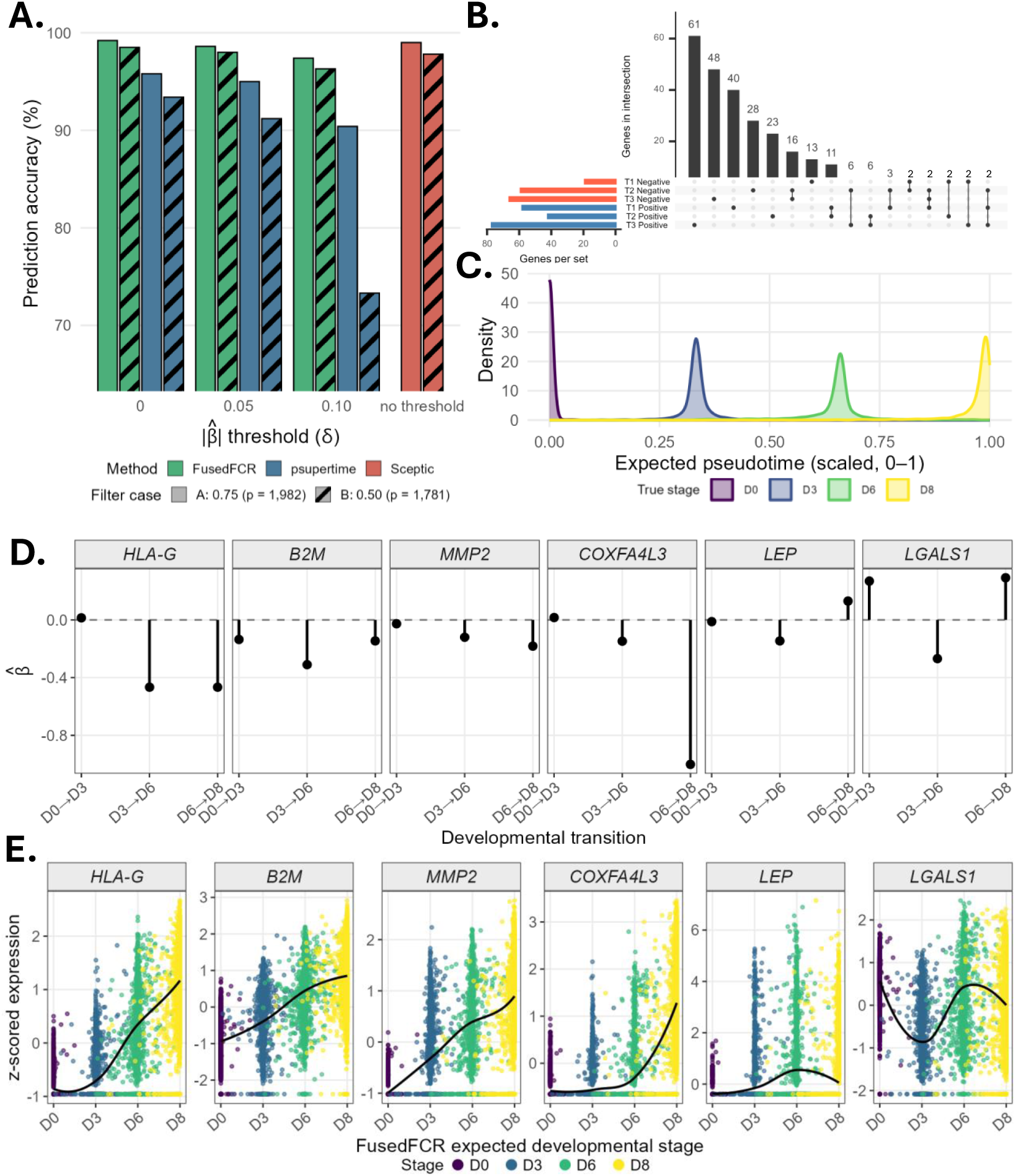
**A.** Gene selection prediction accuracy (%) by method and correlation-filtering threshold in the EVT dataset (GSE307832) on the held-out test set. **B.** UpSet plot of genes selected by FusedFCR across the three EVT differentiation transitions (*|β̂| >* 0.1), colored by coefficient sign (positive, blue; negative, red). **C.** Density plot of FusedFCR pseudotime in the held-out test set (Case A, *δ* = 0.10). **D.** Estimated transition-specific FusedFCR coefficients for six representative genes. **E.** Z-scored log expression for the same genes against FusedFCR expected differentiation day, with LOESS-smoothed trends.

Across the three transitions, 269 unique genes were selected in at least one transition, of which 56 were selected at two or more transitions (Figure 4B) showing that a majority of selected genes were specific to a single transition and direction rather than shared broadly across the developmental time course. The six genes in Figure 4D-E were selected to illustrate distinct modes of transition-specific regulation. *HLA-G*, which promotes maternal-fetal immune tolerance (Zhuang et al., 2021), showed no effect at the first transition but a sustained, monotonically increasing negative effect at the two later transitions (*β̂* = −0.47 at both D3 → D6 and D6 → D8), indicating elevated expression becomes progressively more characteristic of advanced differentiation. *COXFA4L3* was similarly zero at the first transition and modest at the second (*β̂*_2_*_,COXFA4L3_* = −0.15), but carried the single largest-magnitude effect in the dataset at the third transition (*β̂*_3_*_,COXFA4L3_* = −1.00), demonstrating that FusedFCR can detect sharp, transition-specific effects that a single shared coefficient model like psupertime cannot capture. *COXFA4L3* dampens pro-inflammatory signaling important for establishing pregnancy at the maternal-fetal interface (Lee et al., 2021a; Ozarslan et al., 2024). *LGALS1* showed a non-monotonic sign reversal, positive at D0 → D3 (*β̂*_1_*_,LGALS1_* = 0.27) and D6 → D8 (*β̂*_3_*_,LGALS1_* = 0.29) and negative at D3 → D6 (*β̂*_2_*_,LGALS1_* = −0.27); corresponding to an expression dip at intermediate pseudotime before recovery, possibly reflecting its duel hormonal regulation during decidualization as the body prepares the uterine lining for pregnancy (Jovanović Krivokuća et al., 2021). This reversal helps to exemplify that the fusion penalty does not smooth away genuine directional changes in underlying biology, and is able to capture the dynamic nature of gene expression. *B2M* and *MMP2*, key for trophoblast invasion during early implantation (Lee et al., 2019), both showed consistently negative effects across all three transitions, with

*B2M* peaking in magnitude at the second transition (*β̂*_2_*_,B2M_* = −0.31) and *MMP2* strengthening monotonically toward the third (*β̂_,M MP_* _2_ = {0, −0.12, −0.18}), tracking steadily increasing expression for both genes. Finally, *LEP*, which promotes EVT cell invasion in early pregnancy (Wang et al., 2014; Huang et al., 2023), showed a modest sign reversal from zero at the first transition to a small negative effect at second (*β̂*_2_*_,LEP_* = −0.15) to a positive effect at the third (*β̂*_3_*_,LEP_* = 0.13), consistent with expression rising through the intermediate stage before partially declining and then rising again. Together, these genes show how directional shifts and transition-specific effects, which are obscured entirely by a single shared coefficient, emerge naturally from a per-transition model.

Transition and direction-specific GSEA identified significant enrichment among genes selected at the transition, D3 → D6 transition, described in the original study as pivotal to EVT differentiation (Moreno-Irusta et al., 2026). The GO biological process terms female pregnancy (GO:0007565), multi-organism reproductive process (GO:0044703), and multi-multicellular organism process (GO:0044706) were each enriched at FDR-adjusted *p* = 0.0057, driven by the same nine genes: *HPGD*, *FBLN1*, *MMP2*, *LEP*, *PSG6*, *PSG2*, *PSG9*, *PSG5*, and *PSG3*. Both *MMP2* and *LEP*, highlighted above, were among the driving genes, providing convergent coefficient and pathway-level evidence for this transition as a regulatory switch point in EVT differentiation. Together, these results indicate that FusedFCR recovers a biologically coherent, stage specific program of EVT differentiation, capturing hormonal, immune-modulatory, and structural gene expression changes characteristic of the changes required at the maternal-fetal interface to support pregnancy, rather than a single time-invariant gene signature (Huang et al., 2023).

#### 2.2.5 Human Induced Pluripotent Stem Cells Differentiation

Next, we analyzed a human induced pluripotent stem cell (iPSC) derived astrocytes dataset (Yi et al., 2024) collected during astrocyte differentiation over three weeks (D0, D1, D3, D8, D14, D21). There are 22,087 genes and 43,706 cells. Following quality control filtering (Section 2.2.1) *n* = 43,009 cells remained. Case A retained *p* = 1,987 HVGs and Case B retained *p* = 1,801 HVGs. FusedFCR, psupertime, and Sceptic were fir to the training set (80%; *n* = 43,009) and evaluated on the held-out test set (20%; *n* = 8,602).

In Case A, FusedFCR achieved a test-set prediction accuracy of 99.6%, compared with 96.8% for psupertime and 99.0% for Sceptic (Fig. 5A). As with the previous analyses, increasing the coefficient threshold, *δ*, reduced the prediction accuracy modestly relative to the the number of genes selected for FusedFCR and psupertime. The number of uniquely selected genes for FusedFCR decreased by 83.1%, while the prediction accuracy decreased by 10.5%, when increasing *δ* from 0 to 0.1. Similarly, for psupertime the number of selected genes dropped by 89.4%, but experienced a 50.6% reduction in prediction accuracy for the same change in *δ*. In Case B the contrast was even starker where FusedFCR reduced the number of selected genes by 79.6%, with a 9.8% reduction in prediction accuracy while psupertime reduced the selected genes by 87.4% but exhibited a 59.8% reduction in accuracy, which is consistent with the previous analyses.

**Figure 5:**
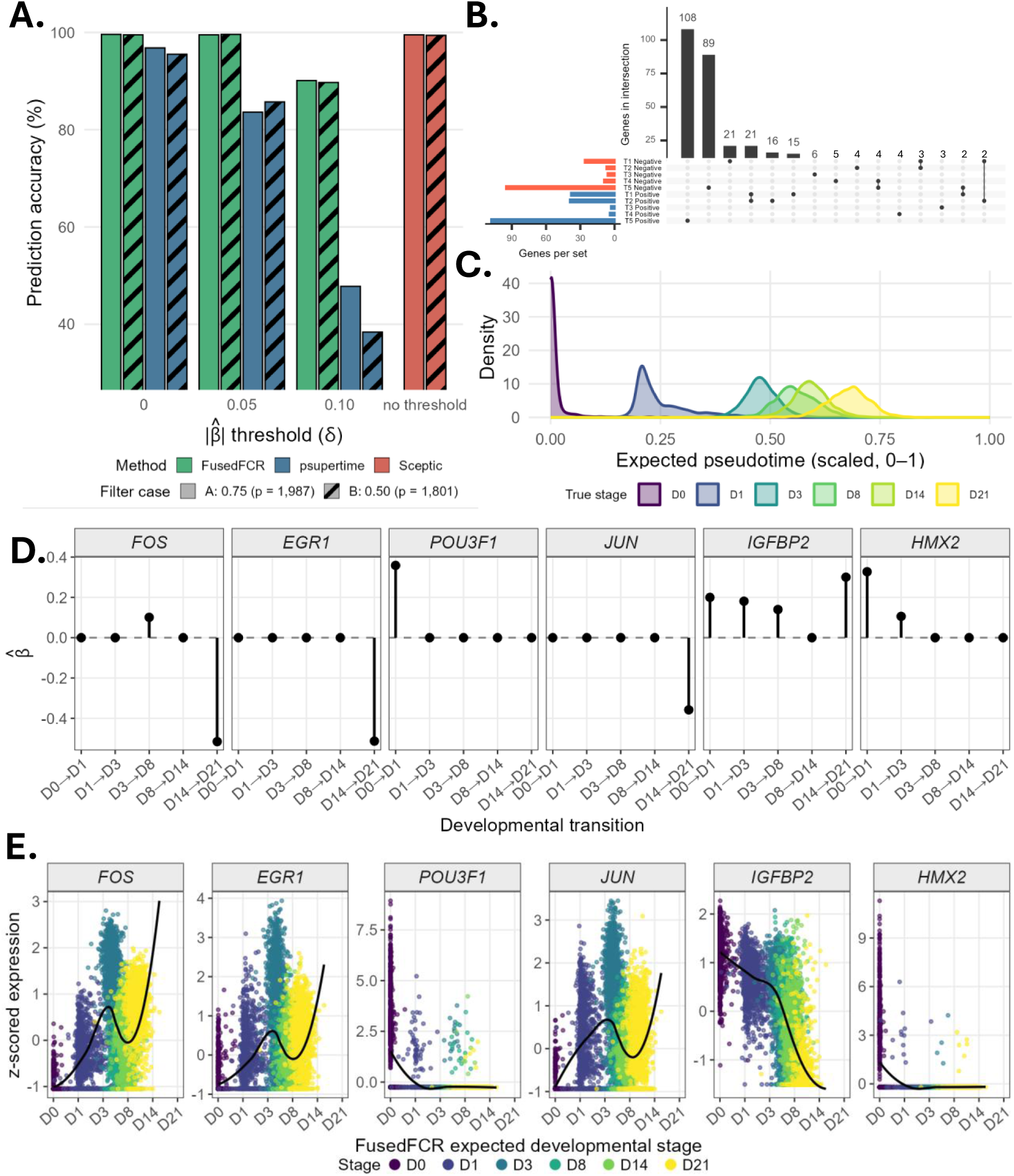
**A.** Gene selection prediction accuracy (%) by method and correlation-filtering threshold in the human iPSC-derived astrocyte dataset (GSE245169) on the held-out test set. **B.** UpSet plot of genes selected by FusedFCR across the five differentiation transitions (*|β̂| >* 0.1), colored by coefficient sign (positive, blue; negative, red). **C.** Density plot of FusedFCR pseudotime in the held-out test set (Case A, *δ* = 0.10). **D.** Estimated transition-specific FusedFCR coefficients for six representative genes. **E.** Z-scored log expression for the same genes against FusedFCR expected differentiation day, with LOESS-smoothed trends.

In FusedFCR Case A (*δ* = 0.1), 318 genes were selected across the transitions, with 271 genes selected at only a single transition and 47 selected in at least two transitions (Fig. 5B). *POU3F1* and *HMX2* were both selected almost exclusively at the earliest transition, D0→D1 (*β*_1_*_,POU3F1_* = 0.36; *β*_1_*_,HMX2_* = 0.33), with smaller residual effects at D1→D3 and negligible contributions thereafter. *POU3F1*, a human-specific pluripotency inducer (Kim et al., 2020), promotes neural fate commitment of pluripotent stem cells (Zhu et al., 2014), consistent with the sharp early expression and rapid decline (Fig. 5E). *HMX2* shows a comparable early peak then monotonic decrease, in line with its role as a homeobox transcription factor active during early neural lineage specification (Wang and Lufkin, 2005). *FOS* and *JUN* both play a role in the proliferation of astrocytes (Green et al., 2026; Cruz-Mendoza et al., 2022). *FOS* was selected at two transitions, with the latter being the largest magnitude effect in the dataset (*β*_3_*_,FOS_* = 0.10; *β*_5_*_,FOS_* = −0.52). The *β* estimate trajectories align with the expression plot (Fig. 5D,E), with the dip shown at D3 aligning with the positive estimate at the D3→D8 transition, and the subsequent large negative effect aligns with the increasing trend at the right of the expression plot. For *JUN*, only the D14→D21 transition exceeded the selection threshold (*β*_5_*_,JUN_* = −0.36), consistent with *JUN* ’s established partnership with *FOS* as a component of the AP-1 complex driving astrocyte proliferation (Green et al., 2026; Cruz-Mendoza et al., 2022). The *β* estimates track the expression plot closely for *JUN*, as expression rises through D3 before declining across D8, D14, and D21, a pattern captured by the large negative effect at the final transition. *EGR1* showed a similar profile to *FOS* and *JUN*, with its only significant effect occurring at the D14→D21 transition (*β*_5_*_,EGR1_* = −0.51), the second-largest magnitude effect in the dataset. *EGR1* is a key component of the transcriptional network regulating gliosis (Beck et al., 2008). Notably, all three genes reach their largest-magnitude effect at the same transition (D14→D21), consistent with their shared classification as immediate early genes (Bahrami and Drabløs, 2016). *IGFBP2* was the most broadly selected gene among this set, exceeding threshold at four of the five transitions (*β*_1_*_,IGFBP2_* = 0.20, *β*_2_*_,IGFBP2_* = 0.18, *β*_3_*_,IGFBP2_* = 0.14, *β*_5_*_,IGFBP2_* = 0.30), with effects that are consistently positive and grow again toward D14→D21. This sustained, multi-transition selection is consistent with *IGFBP2* ’s proposed role as a neurotrophic factor spanning embryonic through adult developmental stages (Khan, 2019), distinguishing it from the isolated spike patterns of the other selected genes. Together the persistence of *IGFBP2* ’s selection and the isolated selection of *FOS*, *EGR1*, *POU3F1*, *JUN*, and *HMX2* illustrate two of the profiles that the fused penalty is designed to distinguish, a sustained biological program are retained as multi-transition trajectories, while more transient signals remain sharply localized rather than being diffused across neighboring transitions. At the D14→D21 transition, direction-specific GSEA showed nominal enrichment for MAPK (hsa04010; FDR *p* = 0.045) and JAK-STAT (hsa04630; FDR *p* = 0.045) signaling, both established astrocyte-differentiation regulators (Li et al., 2012; Stipursky et al., 2012; Ageeva et al., 2024) (full results in Supplemental Material Section S2).

#### 2.2.6 Human Endometrial Cell Differentiation

Lastly, we analyzed a human endometrial cell dataset (Wang et al., 2020) collected during the menstrual cycle (D16, D17, D19, D20, D22, D23, D26 since the onset of the menstrual bleeding) concentrated on the leutial or secretory phase where the endometrium prepares for embryo implantation (Alfer et al., 2020; Choo et al., 2023). There are 33,538 genes and 71,032 cells. Following quality control filtering (Section 2.2.1 *n* = 64,922 cells remained. Case A retained *p* = 1,988 HVGs and Case B retained *p* = 1,735 HVGs. FusedFCR, psupertime, and Sceptic were fit to the training set (80%; *n* = 51,938) and evaluated on the held-out test set (20%; *n* = 12,984).

FusedFCR achieved a test-set prediction accuracy of 97.6%, compared with 53.3% for psupertime and 99.1% for Sceptic, for Case A with *δ* = 0.10 (Fig. 6A). As in the previous analyses, prediction accuracy and the number of genes selected both declined as *δ* increased. However, unlike the other analyses in Case A, FusedFCR and psupertime showed a more comparable drop in prediction accuracy as *δ* increased from 0 to 0.10, an 18.9% decrease for FusedFCR versus 17.4% for psupertime. This was accompanied by a substantial drop in the number of genes selected: FusedFCR’s gene set fell 84.2%, from 1,872 at *δ* = 0 to 314 at *δ* = 0.10, while psupertime’s selected gene count dropped by 97.2%. Despite the comparable relative decline in accuracy, FusedFCR retained superior prediction accuracy across all *δ* thresholds, since its starting accuracy was much higher; psupertime’s accuracy, by contrast, was only slightly better than chance or worse throughout. In Case B, FusedFCR showed a similar pattern to Case A, with a 16.9% drop in accuracy from 96.3% at *δ* = 0 to 80.0% at *δ* = 0.10. psupertime, however, showed a steeper 30.8% drop, from 50.9% to 35.2%, again performing at or below chance level.

**Figure 6:**
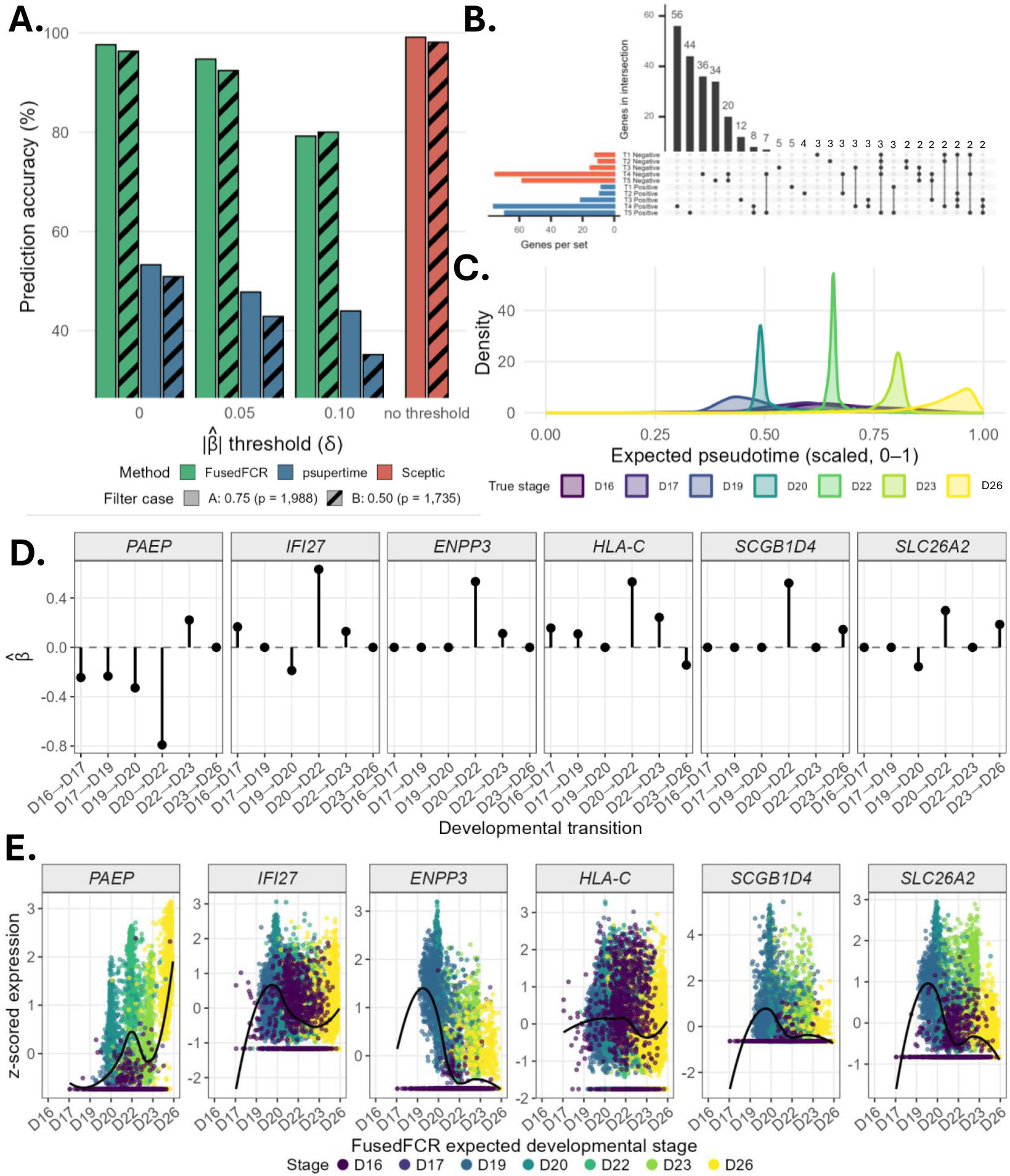
**A.** Gene selection prediction accuracy (%) by method and correlation-filtering threshold in the human endometrium dataset (GSE111976) on the held-out test set. **B.** UpSet plot of genes selected by FusedFCR across the six endometrial transitions (*|β̂| >* 0.1), colored by coefficient sign (positive, blue; negative, red). **C.** Density plot of FusedFCR pseudotime in the held-out test set (Case A, *δ* = 0.10). **D.** Estimated transition-specific FusedFCR coefficients for six representative genes. **E.** Z-scored log expression for the same genes against FusedFCR expected developmental stage, with LOESS-smoothed trends.

Unlike the previous analyses, where 20% or fewer of the selected genes were shared across more than one transition, FusedFCR Case A (*δ* = 0.10) selected 314 genes across the six transitions for the endometrial data, with 209 selected at only a single transition and 105 (33.4%) selected at two or more (Fig. 6B). This dataset is also more complex, with earlier stages having fewer cells than later stages, and the density plot (Fig. 6C) shows overlap among the first three stages but clear separation among the latter four. *PAEP*, which is a key marker of the secretory phase and endometrial differentiation (Jiang et al., 2024), showed a negative effect that strengthened monotonically across the first three transitions, peaked at D20 → D22 transition (*β*_4_*_,PAEP_* = −0.79, the largest magnitude in the dataset), reversed sign to a moderate positive effect at D22 → D23 (*β*_5_*_,PAEP_* = 0.22), and dropped out of selection at the final transition, consistent with the expression trajectory in Figure 6E. *IFI27*, associated with implantation (Khatun et al., 2021), showed a nonmonotonic pattern with a modest positive effect at D16 → D17 (*β*_1_*_,IFI27_* = 0.17), no effect at the second transition, a sign reversal to negative at D19 → D20 (*β*_3_*_,INF27_* = −0.19), the dataset’s second largest effect at D20 → D22 (*β*_4_*_,INF27_* = 0.63), a smaller effect D22 → D23 (*β*_5_*_,INF27_* = 0.13), and dropout at the final transition. This complex effect trajectory, which coincides with the expression plot, illustrates a directional reversal that a single shared-coefficient model like psupertime cannot represent. *ENPP3* and *SCGB1D4* were each selected at only two transitions, both carrying their dominant effect at D20 → D22 (*β*_4_*_,ENPP3_* = 0.53; *β*_4_*_,SCGB1D4_* = 0.52) with a smaller secondary effect at D22 → D23 for *ENPP3* (*β*_5_*_,ENPP3_* = 0.11 and at D23 → D26 for *SCGB1D4* (*β*_6_*_,SCGB1D4_* = 0.14), reflecting a sharp, narrow window of importance rather than gradual accumulation. *HLA-C*, known to be expressed in secretory endometrium and birnds with ILT2 to simulation growth promoting factors (Nilsson and Hviid, 2022), showed modest positive effects at the first two transitions (*β*_1_*_,HLA-C_* = 0.16; *β*_2_*_,HLA-C_* = 0.11), its largest effect at D20 → D22 (*β*_4_*_,HLA-C_* = 0.53), a sustained positive effect at D22 → D23 (*β*_5_*_,HLA-C_* = 0.24), and a late reversal to negative at D23 → D26 (*β*_6_*_,HLA-C_* = −0.14). *SLC26A2*, a marker of the window of implantation (WOI) and important for endometrial receptivity (Kasvandik et al., 2020), was selected at three of the six transitions, initially negative at the third transition from D19 → D20 (*β*_1_*_,SLC26A2_* = −0.16) before sign reversing at D20 → D22 (*β*_4_*_,SLC26A2_* = 0.30), no selection at the fifth transition and a moderate positive effect at D23 → D26 (*β*_6_*_,SLC26A2_* = −0.23), consistent with the transient mid-trajectory expression bump visible in Figure 6E.

The six genes highlighted here each consistently have their largest coefficient magnitudes on the D20 → D22 transition, and additionally for *PAEP*, *IFI27*, and *SLC26A2* their directional reversals. This is particularly notable since this transition corresponds closely to the classically defined WOI, the brief period of maximal endometrial receptivity occurring around days 20–24 of a regular cycle where molecular and structural changes occur that impact the endometrial receptivity for embryo implantation (Alfer et al., 2020; Choo et al., 2023; Luongo et al., 2025). Transition and direction specific GSEA situate these coefficient patterns within this broader biological context. At D16 → D17, up-regulated genes were enriched for type I interferon response (GO:0034340, GO:0060337, GO:0071357). All three type I interferon response pathways were enriched at FDR-adjusted p = 0.0041 and driven by the same three genes: *IFITM3*, *IFITM2*, *IFI27*), directly implicating *IFI27* ’s positive effect at this earliest transition. At D19 → D20, immediately preceding the WOI, down-regulated genes were enriched for antibacterial (GO:0019731; FDR-adjusted p = 0.017) and innate (GO:0002227; FDR-adjusted p = 0.021) immune response (*SLPI*, *DEFB1*, *RPL39*), directly implicating both *PAEP* and *IFI27* in a coordinated down-regulated program that precedes their reversal at D20 → D22. At D20 → D22 itself, up-regulated genes were enriched for the complement and coagulation cascade (hsa04610; *CLU*, *CFD*, *SERPINA5*, *PLAT*, *PLAUR*, *C1S*), a pathway repeatedly identified as one of the most significantly enriched at the implantation site, reflecting endometrial preparation for the remodeling and immune tolerance required for embryo implantation and growth (Bastu et al., 2019). Together, these results indicate that FusedFCR recovers a biologically coherent, stage-specific program of endometrial decidualization, capturing interferon-response, immune-tolerance, and secretory gene expression changes that converge sharply on D20 → D22, the transition mapping onto the clinically established WOI, as the pivotal transcriptional switch point, rather than a single time-invariant gene signature.

## 3 Discussion

In this study, we introduced FusedFCR, a fused lasso-based forward continuation-ratio model for gene selection and prediction in ordinal time-course scRNA-seq data. Rather than assuming that gene effects remain constant throughout the developmental trajectory, FusedFCR estimates gene-specific effects at each transition, allowing it to capture localized, transient, and sign-changing associations. Across simulated and four real datasets, the method achieved accurate stage prediction while recovering transition-specific gene-effect patterns.

This structure reflects the dynamic nature of developmental regulation, in which a gene may be active at one transition, inactive at another, or reverse its direction of association. Shared-effect models such as psupertime (Macnair et al., 2022) instead estimate an average effect across the trajectory, while pseudotime-based marker detection methods generally identify genes associated with a continuous latent ordering without explicitly assigning their effects to observed transitions. In simulations, FusedFCR more accurately recovered the underlying transition-specific gene-effect trajectories and generally achieved higher predictive accuracy. In the beta-cell, EVT, iPSC derived astrocyte, and endometrium datasets, its prediction accuracy remained high as increasingly stringent coefficient thresholds removed genes with smaller estimated effects, indicating that the larger coefficients retained most of the predictive information. In contrast, psupertime showed substantially greater declines in accuracy under the same thresholding scheme, suggesting that many of its smaller coefficients still contributed to prediction. Consequently, coefficient magnitude may be a less reliable basis for gene ranking or for constructing downstream pseudotime projections in psupertime. Although Sceptic (Li et al., 2025) achieved comparable predictive performance in several settings, its lack of interpretable gene-level coefficients limits its utility for transition-specific gene selection and biological interpretation. Together, these results highlight the key advantages of FusedFCR, allowing for transition-specific coefficients that capture dynamic gene effects across transitions, while the lasso and fused penalties jointly guard against overfitting, the model achieves both accurate prediction and stable, interpretable gene-level inference, in a way neither a single shared coefficient model like psupertime not a predictive model like Sceptic can offer.

Our simulations indicated some underestimation of coefficient magnitudes, particularly under sparse-signal settings, and tuning the sparsity and fusion parameters by cross-validation can be computationally demanding. Future work could address these limitations through non-convex penalties such as SCAD or MCP (Liu et al., 2022; Sun and Cui, 2026), which may reduce shrinkage bias, or through a Bayesian formulation using sparsity-inducing priors such as spike-and-slab or horseshoe distributions (Ishwaran and Rao, 2005; Carvalho et al., 2010). A Bayesian extension would also allow regularization parameters and coefficients to be estimated jointly while providing posterior credible intervals and a principled characterization of selection uncertainty beyond post-hoc |*β*| thresholding. The FusedFCR framework could also be extended to accommodate more complex study designs. In multi-sample single-cell studies, sample- or condition-specific effects could be incorporated to separate patient-level heterogeneity from the shared developmental trajectory (Liu et al., 2022; Campbell and Yau, 2016; Hou et al., 2023; Roux de Bézieux et al., 2024). Similarly, applications to spatially resolved time-course data could introduce neighborhood-based penalties or spatial random effects to account for dependence among nearby cells rather than treating them as independent (Patino-Martinez et al., 2026; Seal and Neelon, 2026).

We developed an efficient, user-friendly R package implementing FusedFCR, with functions for model fitting, cross-validated hyperparameter tuning, and visualization of transition-specific coefficients. The package will be made available on GitHub.

## 4 Methods

We consider time-course scRNA-seq datasets in which cells are sampled at known discrete time points, providing population-level snapshots of a dynamic biological process. Supervised pseudo-time inference methods leverage these time labels to learn a refined, continuous ordering of cells along the underlying trajectory. Here, we review two such computational methods for pseudotime inference: psupertime (Macnair et al., 2022) and Sceptic (Li et al., 2025).

### 4.1 Review of psupertime and Sceptic

Let there be *N* cells with *y_i_* ∈ {1*, . . ., T* } denoting the experimental collection time or stage of the *i^th^* cell for *i* = 1*, . . ., N* . Let **x***_i_* = (*x_i_*_1_*, . . ., x_ip_*)^⊤^ ∈ R*^p^* denote its *p*-dimensional transcriptomic profile. We assume a strict chronological ordering among the temporal stages such that 1 *<* 2 *< . . . < T* . psupertime considers two classes of ordinal regression models: the proportional odds (PO) model and the CR model. The PO model captures the log-odds of the cumulative probability as a linear function of the transcriptomic profile:

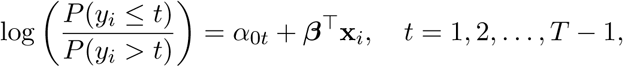

where ***β*** = (*β*_1_*, . . ., β_p_*)^⊤^ ∈ R*^p^* represents the feature-specific effects and *α*_0_*_t_* denotes the stage-specific intercepts. For notational simplicity, we suppress conditioning on **x***_i_* when it is clear from context. Under this parameterization, a large positive value of *β_j_* increases the odds of a cell transitioning to later developmental stages. Alternatively, the CR model focuses directly on sequential state transitions. Specifically, the forward CR or FCR models the discrete hazard—the conditional probability that a cell exits the progression at stage *t*, given that it has survived up to that stage:

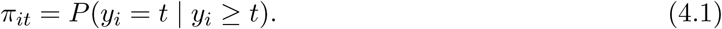

This conditional probability is similarly modeled using a logit link function:

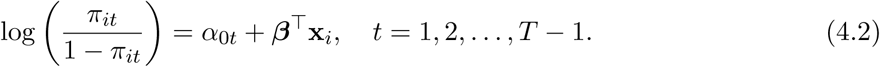

Unlike the PO framework, which measures global shifts across cumulative distributions, the FCR explicitly models the stage-to-stage transition hazard. In this parameterization, ***β*** quantifies the feature’s influence on the probability of a cell progressing to subsequent stages versus terminating at the current stage. By explicitly modeling the sequential transitions, the FCR framework directly mirrors the directional nature of cellular differentiation, providing a structurally superior foundation for elucidating the drivers of developmental progression. To facilitate variable selection, psupertime uses a lasso or *ℓ*_1_ penalty on ***β***, leading to the minimization problem:

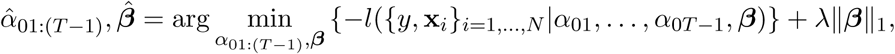

where *l*(·) is the log-likelihood function and ∥***β***∥_1_ denote the *ℓ*_1_ norm of ***β***. Under this regularized framework, any feature retaining a non-zero magnitude (*β̂_j_*= 0) can be interpreted as a crucial driver of the overall cellular trajectory. Finally, psupertime defines pseudotime as the linear projection ***β̂***^⊤^**x***_i_*. This approach is useful for trajectory inference and identifying broad developmental drivers, but it assigns each gene a single coefficient across all time points. This makes the model less suited for detecting stage-specific markers, because a gene that is important at one transition may have little or no effect at another. This is a particular disadvantage in applications where one is interested in capturing developmentally dynamic genes whose expression changes temporally (Cardoso-Moreira et al., 2019).

In contrast to the ordinal modeling approach of psupertime, Sceptic adopts a nominal classification strategy, modeling experimental times as independent discrete classes via one-vs-rest support vector machines (SVMs) (Cristianini and Shawe-Taylor, 2000). Their model estimates the conditional class probability *P* (*y_i_* = *t* | **x***_i_*) for a given profile **x***_i_*, and ultimately defines the cell’s continuous pseudotime through the expected value 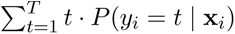. While Sceptic improves predictive accuracy over psupertime for time course prediction, it does not explicitly produce individual gene effects, making it ill-suited for identifying genes that may drive cellular progression. This is a central goal in our case studies, where, for example, recovering the transition specific markers of iPSC astrocyte differentiation (GSE245169; Yi et al. (2024)) requires stage specific individual gene effects. This gap motivates our fused stage-specific coefficient approach which retains predictive competitiveness while directly yielding interpretable, transition-specific gene effects.

### 4.2 Extension to the FusedFCR model

To circumvent the limitations of psupertime, we replace the fixed coefficient vector ***β*** in equation (4.2) with stage-specific coefficients ***β****_t_* = (*β_t_*_1_*, . . ., β_tp_*)^⊤^. The forward continuation probability *π_it_* changes as

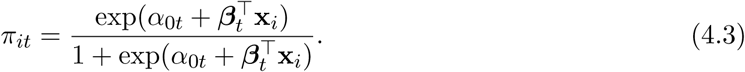

To derive the log-likelihood of the complete data, we need the unconditional probabilities *p_it_* = *P* (*y_i_* = *t*). Utilizing (4.1) and (4.3), *p_it_* can be expressed as

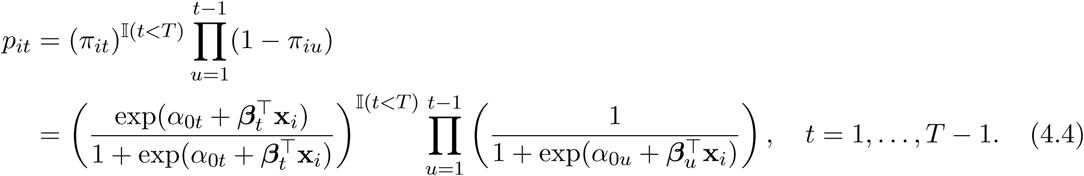

The final-stage probability is determined by the first *T* −1 conditional probabilities as 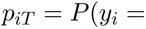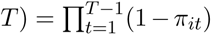, so only (*T* − 1) forward continuation probabilities need to be modeled (Neelon et al., 2019). Crucially, ***β****_t_* quantifies stage-specific covariate effects and can therefore identify genetic drivers that are unique to each transition, a capacity that neither psupertime, which estimates a single gene effect pooled across the entire trajectory, nor Sceptic, which offers no feature effects, is able to provide. This distinction has practical implications in time course scRNA-seq data, where transition-specific coefficients make it possible to characterize a genes effect across developmental transitions, offering insight into the drivers of cellular development or differentiation. To illustrate, consider the first covariate *x_i_*_1_ and its corresponding coefficient *β_t_*_1_ at stage *t*, while holding all other covariates fixed. A positive coefficient, *β_t_*_1_ *>* 0, indicates that increasing *x_i_*_1_ raises the conditional probability of exiting the developmental trajectory at stage *t*, given that the cell has reached that stage. Conversely, a negative coefficient, *β_t_*_1_ *<* 0, lowers this probability and increases the conditional probability that the cell continues to a subsequent stage. Complete derivations are provided in Section S3, Supplementary Methods.

Using equation (4.4), we can express the joint data likelihood as,

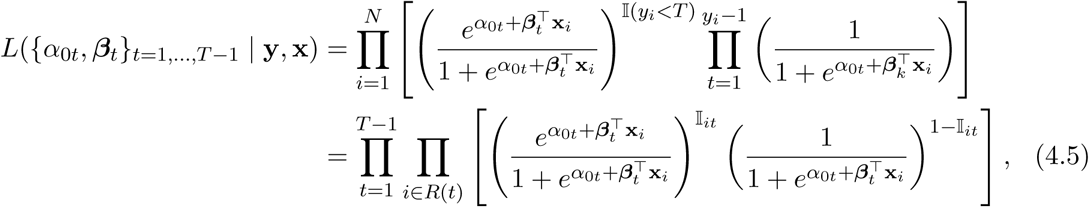

where Ι*_it_* = Ι(*y_i_* = *t*) and *R*(*t*) = {*i* : *y_i_* ≥ *t*}. Equation (4.5) contains a total of (*T* − 1)(*p* + 1) parameters. To perform feature selection, we impose an *ℓ*_1_ penalty on the stage-specific coefficients ***β****_t_*. Because biologically active features may exhibit similar effects across adjacent stages, we additionally penalize differences between neighboring coefficients. In practice, this prevents overfitting to the noise of a single transition and instead leverages the greater context of the sequential process that time-course scRNA-seq data captures, reflecting the correlated nature of cellular development across adjacent stages (Cardoso-Moreira et al., 2019). The resulting fused lasso penalty is

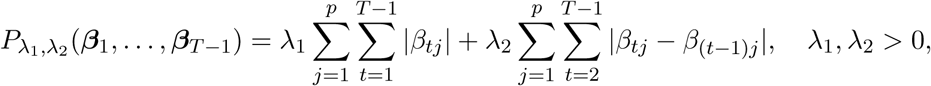

where |·| denotes the absolute value of a variable. Let 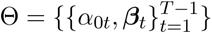 denote the full parameter set, and let *l*(Θ | **y**, **x**) = log *L*(Θ | **y**, **x**) denote the corresponding log-likelihood. Finally, we estimate Θ using the penalized loss function:

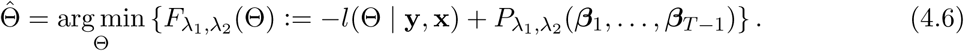

The tuning parameter *λ*_1_ *>* 0 controls the overall sparsity of the stage-specific coefficients, whereas *λ*_2_ ≥ 0 controls the degree of similarity between coefficients at adjacent stages. The stage-specific intercepts *α*_0_*_t_* are left unpenalized, allowing the baseline transition probabilities to vary freely across stages. Parameter estimation is described in Section 4.3. After the parameters are optimized, we define pseudotime for FusedFCR as pseudotime 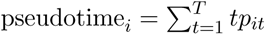 similar to Sceptic.

### 4.3 Parameter estimation and hyperparameter selection

To facilitate parameter estimation, the FCR log-likelihood *l*(Θ | **y**, **x**) can be represented in standard generalized linear model (GLM) form. Utilizing the binary indicator variables Ι*_it_* for each individual *i* across their surviving stages up to *y_i_*, the joint log-likelihood algebraically decomposes into a sum of independent Bernoulli log-likelihoods:

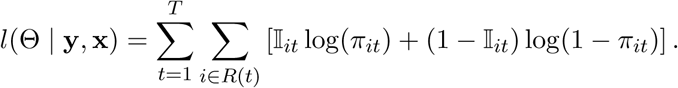

Since this structure mirrors a standard logistic regression over the expanded pseudo-dataset {Ι*_it_* | *i* = 1*, . . ., N, t* = 1*, . . ., y_i_*}, we can utilize the standard Newton-Raphson scoring method to optimize the unpenalized stage-specific intercepts, 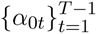 (Nelder and Wedderburn, 1972; Wedderburn, 1974; Cox, 1984) from equation (4.6). The non-differentiability of the fused lasso penalty precludes applying the same approach to 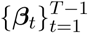. We therefore consider the dynamic programming algorithm of Johnson et al. (2013) (Johnson, 2013) to optimize the penalized coefficients. Specifically, the algorithm first sets *λ*_1_ = 0 and solves the fused lasso signal approximator (FLSA) problem for a fixed *λ*_2_ *>* 0. For each feature *j*, it updates the stage-specific coefficient sequence 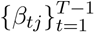 to obtain intermediate estimates 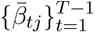 under the fusion penalty alone. The sparsity penalty is then incorporated through soft-thresholding, yielding *β̂_tj_*= sign(*β̂_tj_*) max {0, |*β̂_tj_*| − *λ*_1_} (Friedman et al., 2007). The procedure is summarized in Algorithm 1, with more details on the overall numerical stability of the optimization provided in the Supplementary Methods, Section S4. A similar strategy was applied by Adhikari et al. (2019) to multinomial logistic regression with a fused lasso penalty.

The sparsity and temporal smoothness of the model are controlled by *λ*_1_ and *λ*_2_, respectively. Large values of *λ*_1_ may remove important covariates, whereas large values of *λ*_2_ may oversmooth

#### Algorithm 1: Parameter estimation for FusedFCR model parameters

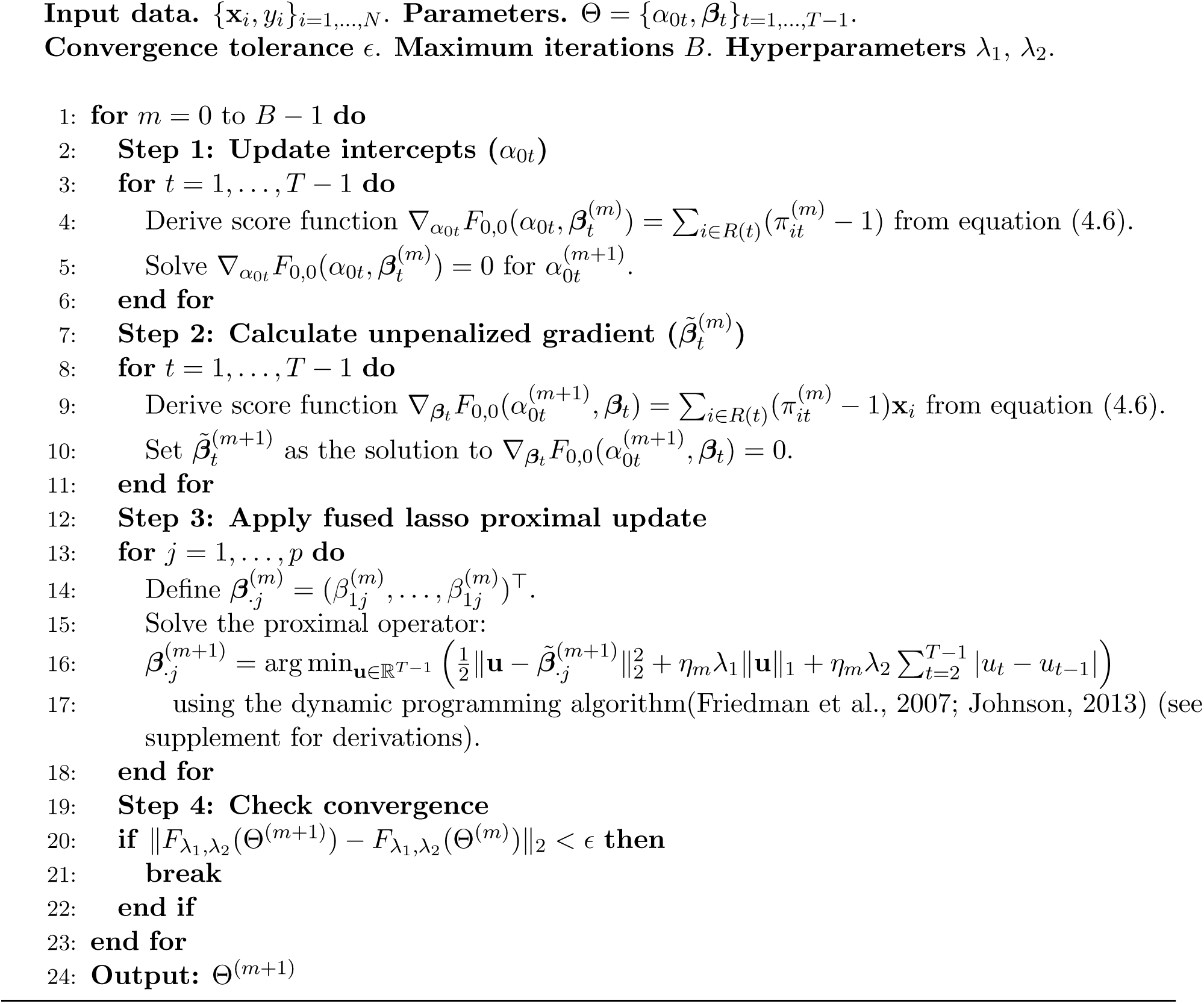

stage-specific effects. Conversely, values that are too small can produce dense models that obscure transition-specific genetic signals. We therefore select *λ*_1_ and *λ*_2_ in a data-driven manner using multi-fold cross-validation (CV), following established practice in related penalized regression settings (Tibshirani et al., 2005; Adhikari et al., 2019; Xin et al., 2014; Macnair et al., 2022). Standard random CV can produce imbalanced training and validation folds when cells are unevenly distributed across developmental stages. To improve stability, we therefore use repeated stratified CV, preserving the stage distribution within each fold (Purushotham and Tripathy, 2011; Berrar et al., 2019; Szeghalmy and Fazekas, 2023). We select the pair (*λ*_1_*, λ*_2_) that minimizes the average cross-entropy loss across repetitions. We also considered the Extended Bayesian Information Criterion (EBIC), a widely used criterion for variable selection in high-dimensional settings (Chen and Chen, 2012; Mallick and Yi, 2013; Yue et al., 2026). Although EBIC produced stable feature subsets, it substantially reduced out-of-sample predictive performance in both simulated and real scRNA-seq datasets. Because our objective is to balance feature selection with predictive accuracy, we primarily focus on the CV-based tuning procedure. Fig. 7 gives a graphical summary of FusedFCR.

**Figure 7:**
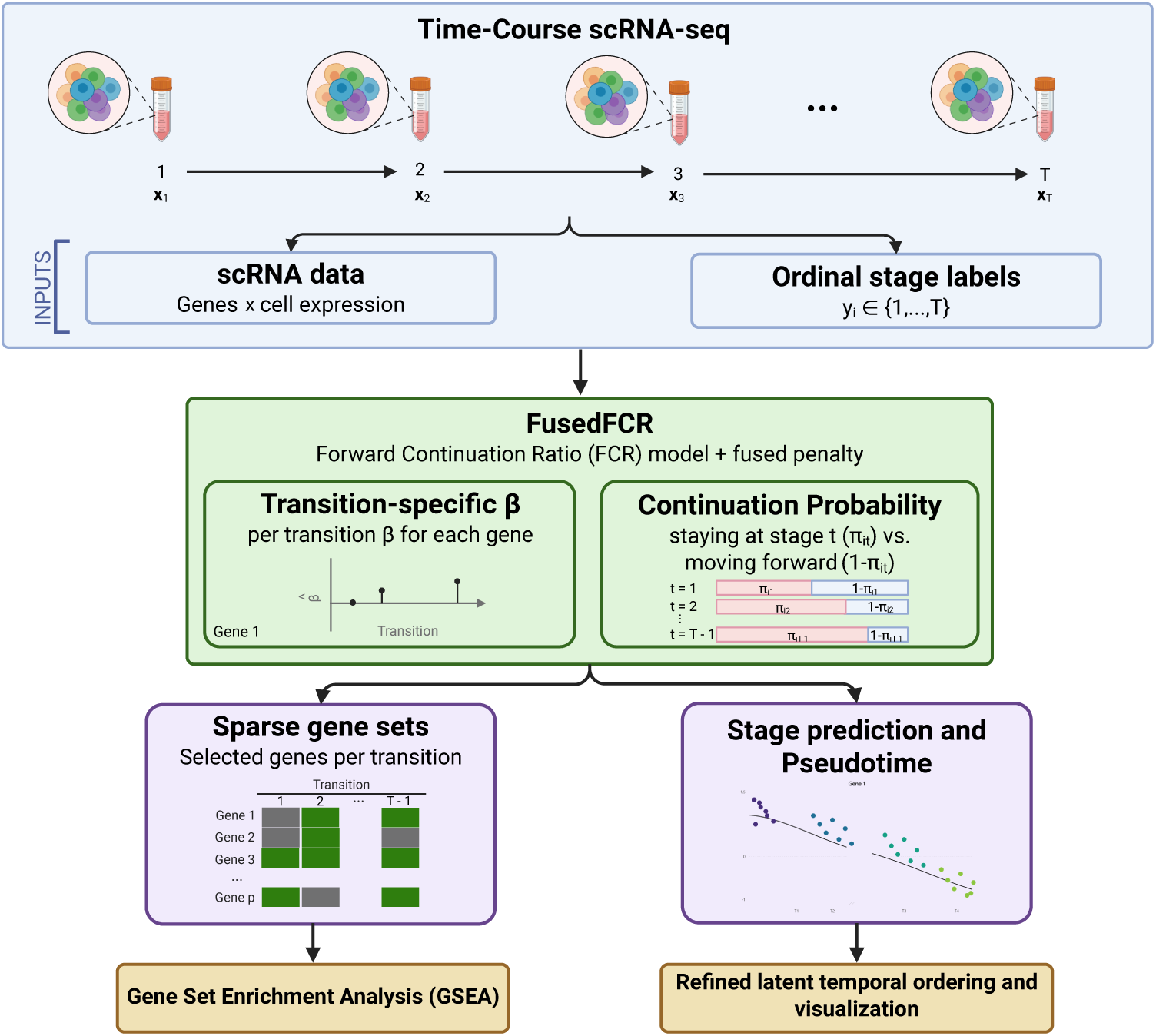
Graphical summary of FusedFCR. Created with BioRender (https://biorender.com/).

## 5 Data Access

The scRNA-seq data used in this study are available through the NCBI Gene Expression Omnibus with accession numbers GSE87375 (mouse embryonic beta cells), GSE307832 (human EVT cells), GSE245169 (human iPSC), and GSE111976 (human endometrium). The scRNA-seq data for EVT differentiation was downloaded from https://github.com/genlab/2026-evt_public.git.

## Acknowledgments

C.M., A.C., B.N., E.H., and S.S. were supported in part by the Data Science Shared Resource, Hollings Cancer Center, Medical University of South Carolina (P30 CA138313). S.S. and P.A. were supported by NIH R21 CA28628701A1. S.S. and A.C were supported by the American Cancer Society Institutional Research Grant: IRG-24-129055323-IRG. The content is solely the responsibility of the authors and does not necessarily represent the official views of the American Cancer Society, the National Cancer Institute, and the National Institutes of Health.

