## Supplemental Material for "FusedFCR: A Fused Forward Continuation-Ratio model for marker selection along cell-fate trajectories"

#### S1 Simulation Details

An important practical consideration in simulating from the FCR model is the relationship between the intercept vector $\boldsymbol{\alpha}_t$, the true $\beta$ coefficients, the resulting label distribution, and prediction accuracy under the true model. In the non-temporal scenario, $\boldsymbol{\alpha}_0$ was chosen to

†These authors contributed equally to this work.

produce approximately balanced observation counts across the six outcome categories; however, even under the true model, prediction accuracy was often below 50%, likely reflecting a combination of unstructured predictors and overlapping label distributions that make it difficult to recover the true label. In the temporal scenario, $\boldsymbol{\alpha}_0$ was instead chosen to yield high true-model accuracy, at the cost of label imbalance, to better reflecting diseases with either a high rate of early detection/curability or high mortality. Given these differences, we adopt distinct primary evaluation criteria for each scenario: for the non-temporal scenario, where true-model accuracy is low, we focus on each method's ability to recover the true coefficients across transitions; for the temporal scenario, where prediction is well-defined under the true model, we emphasize predictive accuracy.

##### S1.1 Temporal signal structure

For the temporal scenario, structured signals were added for the first three genes as smooth functions of the underlying developmental time $u_i$, with Gaussian background variation added:

$$\begin{aligned}
X_{i1} &= X_{i1}^{(0)} + 4.5\exp(-5.5u_i),\\
X_{i2} &= X_{i2}^{(0)} + 5\exp\left(-\frac{(u_i-0.5)^2}{(2*0.11^2)}\right),\\
X_{i3} &= X_{i3}^{(0)} + 4.5\exp(-5.5(1-u_i)).
\end{aligned}$$

#### S2 Extended gene set enrichment analysis results

##### S2.1 Embryonic Beta Cells

Transition- and direction-specific gene set enrichment analysis (GSEA), based on KEGG[1–3] and Gene Ontology (GO) biological process terms[4,5] implemented in ShinyGO v0.85.1[6], identified cAMP-mediated signaling as the predominant positively enriched program at the first transition. The strongest KEGG enrichment was observed for the cAMP signaling pathway (mmu04024; FDR $p = 7.99 \times 10^{-5}$), driven by *Npy*, *Crhr2*, *Gip*, and *Sox9*. This finding was further supported by GO biological process terms for cAMP-mediated signaling (GO:0019933; FDR $p = 9.13 \times 10^{-3}$) and its positive regulation (GO:0043950; FDR $p = 2.54 \times 10^{-2}$). In pancreatic beta cells, cAMP functions as a central intracellular second messenger involved in cell fate, survival, proliferation, and differentiation[7].

##### S2.2 Human Induced Pluripotent Stem Cell Differentiation

At the D14→D21 transition, direction-specific GSEA revealed that the MAPK signaling pathway (hsa04010; FDR-adjusted $p = 0.045$) was nominally significant, consistent with the known role of MAPK signaling in glial/astrocyte development and differentiation[8,9]. This pathway was driven by *HSPA1A*, *HSPA8*, *SOCS2*, and *SOCS3*. Overall, the absence of enrichment among the remaining selected genes suggests that many act through effects not well captured by curated GO/KEGG gene sets at current FDR thresholds, consistent with their role as single transcription factors or immediate-early responders rather than members of a broader annotated program. Additionally, at this transition, the JAK-STAT signaling pathway was nominally significant (hsa04630; FDR-adjusted $p = 0.045$) and is known to be important for astrocyte differentiation[10].

#### S3 Derivation of the unconditional probabilities

We start with the definition of $\pi_{it}$ which is given by,

$$\pi_{it} = P(Y_i = t \mid Y_i \geq t) = 1 - P(Y_i > t | Y_i \geq t). \tag{S3.1}$$

In addition, we note that for any set $A$ that is a subset of $B$, i.e., $A \subset B$, we can always write

$$P(A) = P(A \cap B),$$

since both of $A$ and $B$ occur simultaneously only when $A$ occurs. Using the above identity and applying equation (S3.1) recursion, we can derive the unconditional probability $P(Y_i = t)\ \forall t < T$ as,

$$
\begin{aligned}
P(Y_i = t) &= P(Y_i = t, Y_i \geq t) \\
&= P(Y_i = t | Y_i \geq t) P(Y_i \geq t) \\
&= P(Y_i = t | Y_i \geq t) P(Y_i \geq t | Y_i \geq t-1) P(Y_i \geq t-1) \\
&= P(Y_i = t | Y_i \geq t)(1 - P(Y_i = t-1 | Y_i \geq t-1)) P(Y_i \geq t-1) \\
&= \pi_{it}(1 - \pi_{it-1}) P(Y_i \geq t-1) \\
&= \pi_{it}(1 - \pi_{it-1}) P(Y_i \geq t-1 | Y_i \geq t-2) P(Y_i \geq t-2) \\
&= \pi_{it}(1 - \pi_{it-1})(1 - \pi_{it-2}) P(Y_i \geq t-2) \\
&\ \vdots \\
&= \pi_{it}(1 - \pi_{it-1})(1 - \pi_{it-2}) \dots P(Y_i \geq 1) \\
&= \pi_{it} \prod_{u=1}^{t-1} (1 - \pi_{iu}),
\end{aligned} \tag{S3.2}
$$

since $\pi_{i1} = P(Y_i = 1) = 1 - P(Y_i \geq 1)$. Using the notation $p_{it} = P(Y_i = t)$ from Section 4.2, we can write,

$$
\begin{aligned}
p_{i1} + p_{i2} &= \pi_{i1} + \pi_{i2}(1 - \pi_{i1}) = 1 - (1 - \pi_{i1})(1 - \pi_{i2}) \\
p_{i1} + p_{i2} + p_{i3} &= 1 - (1 - \pi_{i1})(1 - \pi_{i2}) + \pi_{i3} \prod_{u=1}^{2} (1 - \pi_{iu}) = 1 - \prod_{u=1}^{3} (1 - \pi_{iu}) \\
&\ \vdots \\
\sum_{u=1}^{T-1} p_{it} &= 1 - \prod_{u=1}^{T-1} (1 - \pi_{iu}).
\end{aligned}
$$

Using the above identities, it is straightforward to show that,

$$P(Y_i = T) = 1 - \sum_{u=1}^{T-1} p_{it} = \prod_{u=1}^{T-1} (1 - \pi_{iu}). \tag{S3.3}$$

Combining equations (S3.2) and (S3.3), we can express $p_{it}$ as,

$$p_{it} = (\pi_{it})^{\mathbb{I}(t<T)} \prod_{u=1}^{t} (1 - \pi_{iu}).$$

### S4 Numerical considerations

#### S4.1 Stabilizing optimization and hyperparameter tuning

The performance of cross validation strategy outlined in Section 4.3 of the main manuscript depends on the number of observations in specific stages. To illustrate this let us define $n_k = \sum_{i=1}^{N} 1\{y_i = t\}$, i.e., the number of observations recorded at a specific stage $t$. In many biological applications, observations vary a lot across stages, and hence $n_1$ may be significantly different from $n_2$. As Adhikari et al. (2019)[11] have observed, when this happens, the objective function outlined in equation 2.6 of the main manuscript performs poorly in terms of prediction as well as feature selection. To avoid this numerical collapsing, we first modify the log-likelihood contribution as,

$$l^a(\{\alpha_{0k}, \boldsymbol{\beta}_k\}_{t=1,\ldots,T} | \mathbf{y}) = \sum_{t=1}^{T} \frac{1}{n_k} \sum_{i:i \in R(t)} \left[ \mathbb{I}_{it}(\alpha_{0k} + \boldsymbol{\beta}_k^T \mathbf{x}_i) - \log\left(1 + e^{\alpha_{0k} + \boldsymbol{\beta}_k^T \mathbf{x}_i}\right) \right].$$

For each of the $T - 1$ stages, we standardize the stage-specific total likelihood contribution by the number of recorded observations in that stage. This mitigates the effect of class imbalance. Consequently, we proceed with the new objective function $F^a_{\lambda_1,\lambda_2}(\Theta)$ which can be expressed as,

$$F^a_{\lambda_1,\lambda_2}(\Theta) = -l^a(\Theta \mid \mathbf{z}) + P_{\lambda_1,\lambda_2}(\boldsymbol{\beta}_1, \ldots .\boldsymbol{\beta}_p) \tag{S4.1}$$

We proceed with the new fused lasso objective (S4.1) for the subsequent parameter estimation steps of $\alpha_{0k}$ and $\boldsymbol{\beta}_k$'s.

##### S4.2 Selection of the optimal learning rate

The score functions in Algorithm 1 in the main manuscript do not have a closed form. A Newtons first order method is generally used in optimizing $\alpha_{0k}$ and $\boldsymbol{\beta}_k$'s, i.e.,

$$
\begin{aligned}
\alpha_{0k}^{(m+1)} &= \alpha_{0k}^{(m)} - \eta_m \nabla_{\alpha_{0m}} F^a_{\lambda_1,\lambda_2}(\Theta^{(m)}) \\
\tilde{\boldsymbol{\beta}}_t^{(m+1)} &= \boldsymbol{\beta}_1^{(m)} - \eta_m \nabla_{\boldsymbol{\beta}_t} l^a(\Theta^{(m)}) \\
\boldsymbol{\beta}_{\cdot j}^{(m+1)} &= \text{prox}_{\lambda_1,\lambda_2,\eta_m}\left(\tilde{\boldsymbol{\beta}}_{\cdot j}^{(m+1)} - \eta_m \nabla_{\boldsymbol{\beta}_{\cdot j}} l^a(\Theta^{(m)})\right) \quad \forall j = 1,\ldots,p,
\end{aligned}
$$

where $\text{prox}_{\lambda_1,\lambda_2,\eta_m}(\cdot)$ is the proximal update. All of these optimization steps depend on the learning rate $\eta_m$ at the $m^{th}$ iteration. We use an backtracking line search algorithm to select $\eta_m$ in a data driven manner. Specifically, at each iteration $m$, we initialize $\eta = \eta_{m-1}$ and repeatedly shrink it by setting $\eta \leftarrow \rho\eta$ until the following condition is satisfied:

$$
l^a(\Theta^+) \leq l^a(\Theta^{(m)}) + \langle \Theta^+ - \Theta^{(m)}, \nabla l^a(\Theta^{(m)}) \rangle + \frac{1}{2\eta} \|\Theta^+ - \Theta^{(m)}\|_2^2,
$$

where $\Theta^+ = \text{prox}_{\lambda_1,\lambda_2,\eta}\left(\Theta^{(m)} - \eta \nabla l^a(\Theta^{(m)})\right)$ is the candidate update. For a more detailed discussion on the backtracking line search algorithm, we refer the authors to Adhikari et al.[11].

[2] M. Kanehisa, M. Furumichi, M. Tanabe, Y. Sato, and K. Morishima. KEGG:

new perspectives on genomes, pathways, diseases and drugs. *Nucleic acids research*, 45(D1):D353–D361, 2017.

[3] M. Kanehisa, M. Furumichi, Y. Sato, M. Kawashima, and M. Ishiguro-Watanabe. KEGG for taxonomy-based analysis of pathways and genomes. *Nucleic acids research*, 51(D1):D587–D592, 2023.

[4] M. Ashburner, C. A. Ball, J. A. Blake, D. Botstein, H. Butler, J. M. Cherry, A. P. Davis, K. Dolinski, S. S. Dwight, J. T. Eppig, et al. Gene ontology: tool for the unification of biology. *Nature genetics*, 25(1):25–29, 2000.

[5] The Gene Ontology resource: enriching a GOld mine. *Nucleic acids research*, 49(D1):D325–D334, 2021.
